# Chemo-Profiling by UPLC-QTOF-MS, GC-MS/MS analysis and *In Vitro* Bioactivity Assessment of *Desmodium gangeticum* DC

**DOI:** 10.1101/2024.10.08.617169

**Authors:** Mamata A. Jagtap, Aditya B. Magdum, Vishwajit V. Lugade, Pratik S. Kamble, Dayanand P. Jayannawar, Mansingraj S. Nimbalkar, Swaroopa A. Patil

**Affiliations:** Department of Botany, Shivaji University, Kolhapur, India

**Keywords:** *Desmodium gangeticum*, Dashmoola, Antioxidant, UPLC-QTOF-MS, GC-MS/MS, Anti-inflammatory, Antibacterial

## Abstract

*Desmodium gangeticum* DC., commonly known as Shalparni, and a member of the Fabaceae family, is a small shrub native to tropical and subtropical regions. The active principles of this plant are primarily concentrated in its roots, which are traditionally used in various medicinal formulations. This study aimed to identify alternative sources of raw material by evaluating the biochemical analysis and bioactivity profiling of both fresh and dried plant parts—roots, leaves, and stems. Extracts were prepared using methanol, ethanol, and distilled water and subjected to phytochemical, antioxidant, anti-inflammatory (HRBC membrane stabilization), and antibacterial assays. Additionally, chemical profiling was performed using FTIR, GC-MS/MS, and UPLC-Q-TOF-MS. This is the first report on *D. gangeticum* as one of the Dashmoola plants for its biochemical analysis, bioactivity, and chemoprofiling of the entire plant. Results indicated that the GC-MS/MS analysis identified 25 volatile compounds in leaves, whereas the roots contained 21. These compounds included fatty acids, ketones, hydrocarbons, alcohols, and esters. UPLC-Q-TOF-MS analysis revealed a diverse array of bioactive compounds: 287 in aqueous leaf extracts and 192 in aqueous root extracts, many of which were identified as bioactive agents. Notably, the leaf extracts exhibited a higher number of bioactive compounds compared to the roots, highlighting the plant’s diverse bioactive compounds in different tissues. The study demonstrates that both the leaves and roots of *D. gangeticum* contain significant therapeutic compounds. Therefore, the leaves could serve as a sustainable alternative to roots as raw material for medicinal preparations, contributing to the conservation of this plant in its natural habitat.

## 1. Introduction

Ayurveda, an indigenous system of traditional medicine, has been practiced for thousands of years. It focuses primarily on plant-based medicines for managing various diseases (Randive et al.,, 2014). Within the Ayurvedic tradition, medicinal plants play a foundational role in healthcare, embodying principles of holistic wellness and reliance on natural therapeutics. Research into Ayurvedic medicinal plants is gaining global attention, with pharmaceutical companies investing in drug discovery from these sources (Kareem and Yoganandham, 2022). In India, herbal medicine, derived from natural sources and particularly medicinal plants, serves as a prevalent form of healthcare. Herbal remedies encompass a range of phytochemical constituents, which are intricate organic compounds inherent to plant species and exhibit diverse biological functionalities. Leveraging their multifaceted biological activities, phytochemicals are harnessed in medicinal contexts as essential therapeutic agents (Jayakumari, 2020). Moreover, phytochemicals play a crucial role as bioenhancers in contemporary pharmacotherapy enhancing the therapeutic efficacy of pharmaceutical agents. In response to challenges such as drug resistance and adverse effects, exploring secondary metabolites derived from medicinal flora offers a promising alternative to synthetic pharmaceuticals. Antioxidant substances, whether natural or synthetic, have various effects on the body. Plants may synthesize antioxidants or develop new antioxidation mechanisms through biochemicals. These molecules are secondary metabolites (Gadade and Patil, 2019). A comprehensive understanding of the functionalities and applications of phytochemicals and secondary metabolites is imperative for advancing phototherapeutics, plant biotechnology, and pharmaceutical development.

Plant-derived antimicrobial compounds hold significant therapeutic promise for combating multidrug-resistant bacterial strains while mitigating the adverse effects commonly associated with synthetic antimicrobial agents. This creates an urgent need to explore novel sources of antimicrobial compounds, especially those derived from medicinal plants. Herbal medications are formulated in various forms such as tablets, powders, and decoctions. Dashmoola occupies a central position in Ayurveda for balancing the three doshas (Vata, Pitta, and Kapha), which represent different bodily energies. Dashmoola comprises a blend of ten plant roots and is renowned for its efficacy in balancing the Tridoshas (Vata, Pitta, and Kapha). Notably, *Desmodium gangeticum* is an important constituent of Dashmoola and is esteemed for its profound medicinal properties (Joshi et al., 2023).

*Desmodium gangeticum* DC. (also known as *Pleurolobus gangeticus* (L.) J. St.-Hil. ex H. Ohashi and K. Ohashi; Family: Fabaceae) is a small shrub found in tropical and subtropical regions. It is sometimes referred to as “Salpan,” “Salpani” in Hindi, and “Shalparni” in Sanskrit. It is commonly used alone or in conjunction with other medications in Ayurvedic, Siddha, and Unani medical systems. It is employed to treat inflammatory disorders of the chest and other conditions caused by “vata” disorders. The plant is known to be a bitter tonic, febrifuge, expectorant, digestive, anti-catarrhal, and antiemetic. The roots have been used for snakebites and scorpion stings (Chopra et al., 1956; Nadkarni, 1976). Reports indicate that the plant possesses hepatoprotective, anti-amnesic, anti-inflammatory, antinociceptive, and wound-healing properties (Prasad et al., 2005; Ghosh et al.,1983; Jain et al., 2006). Chemical investigations have identified alkaloids in *D. gangeticum* with anticholinesterase activity, as well as effects on smooth muscle and central nervous system stimulation (Ghosal et al., 1969; 1972). Additionally, pterocarpanoids present in the plant have been associated with anti-diabetic, anti-ulcer, and anti-fertility properties (Hanumanthachar et al., 2005; Latha et al., 1997). Jadeja and Nakar (2010) proposed ethnomedicinal practices including the oral administration of P. gangeticus root three times daily for managing snake bites.

Several renowned Ayurvedic formulations, such as Dasmularistha, Dasmulakwath, Chitrak Haritika, Dasmoola Kadha, Brahma Rasayana, Dashmoola ark, Dashmoolataila, and Dhanvantratailum, prominently feature the roots of *D. gangeticum*. Due to its alkaloids, vitamins, oils, and mineral content (e.g., calcium, phosphorus, magnesium), traditional Ayurvedic and Unani practitioners use *D. gangeticum* to treat various ailments such as cataracts, typhoid fever, dysentery, bronchitis, asthma, piles, and thyroid disorders (Kirtikar and Basu, 1987; Kumar et al.,2014; Sagar et al., 2011; Akram et al., 2014; Velavan et al., 2012). The therapeutic efficacy of *D. gangeticum* is attributed to its diverse phytochemical profile, particularly its alkaloid components, which confer anti-inflammatory, analgesic, and antioxidant properties. Additionally, it exhibits antioxidant effects against reperfusion injury. The dried roots of the plant demonstrate anti-asthmatic properties across chloroform, water, and ethanolic extracts (Vedpal et al., 2016). Vijayalakshmi et al., (2011) studied the antioxidant and antileishmanial activities, alongside the presence of gangetin, a pterocarpene glycoside.

Understanding the phytochemical composition and bioactivity profiling of plants is crucial for identifying potential sources of novel therapeutic agents. Despite extensive research on the roots of *D. gangeticum*, other plant parts, such as leaves and stems, remain underexplored. There is a need for comprehensive profiling to uncover their potential benefits. This study aims to fill this gap by conducting a comprehensive analysis of *D. gangeticum*. The current investigation assesses the total phenolic, flavonoid, and alkaloid content, along with bioactivity profiling including in vitro antioxidant, anti-inflammatory, and antibacterial activities. The antibacterial activity of *D. gangeticum* leaves, stems, and roots against foodborne pathogens, notably Staphylococcus aureus and Escherichia coli, was evaluated. The biomolecules were identified using GC-MS/MS and UPLC-Q-TOF-MS analyses.

## 2. Material and methods

### 2.1 Materials

Folin-Ciocaltue reagent, Na_2_CO_3_, 2% methanolic AlCl_3_, 0.05M FeCl_3_, 0.05M HCL, 0.05M 1,10-phenanthroline, 300mM sodium acetate buffer, 10mM 2,4,6-Tripyridyl-S-triazine (TPTZ) solution, 20mM FeCl_3_.6H_2_O solution), 0.6 M sulphuric acid, 28mM sodium phosphate, 4mM ammonium molybdate, 2mM FeSO_4_, 5mM ferrozine, Nitroblue tetrazolium (NBT), dimethyl sulfoxide (DMSO), 5mM NaOH, 7mM ABTS, 2.45mM potassium persulfate, 25mM methanolic DPPH (2, 2-diphenyl-1-picrylhydrazyl), Mueller Hinton Agar, Luria Bertani Broth, Methanol, Ethanol, Distilled water, Catechol, Rutin, Colchicine, Ascorbic acid, EDTA, NaCl, sodium phosphate buffer, acetylsalicylic acid (ASA), HRBC, Streptomycin, Microbes *Staphylococcus aureus* (NICM 2654) and *Escherichia coli* (NICM 2832).

### 2.2 Collection of plant material

*Desmodium gangeticum* (L.) DC. was collected from Hiwarkheda (Gautala), Dist. Chhatrapati Sambhajinagar, Maharashtra, India. The plant specimen was identified and processed as dry and fresh. Drying was carried out at 50°C, while fresh materials were stored at -80°C for further analysis.

### 2.3 Preparation of extracts

Fresh and dried leaves, stems, and roots of *D. gangeticum* were used to study biochemical, antioxidant, and antimicrobial activity. The plant extracts (1%) were prepared using overnight soaking, followed by sonication at 50 Hz for 30 minutes (Fig. 1). Three different solvents methanol, ethanol, and distilled water were used for the extraction. Extracts were filtered through Whatman filter paper No.1. and stored in a Borosilicate bottle and kept at -20℃ until used.

**Fig. 1.**
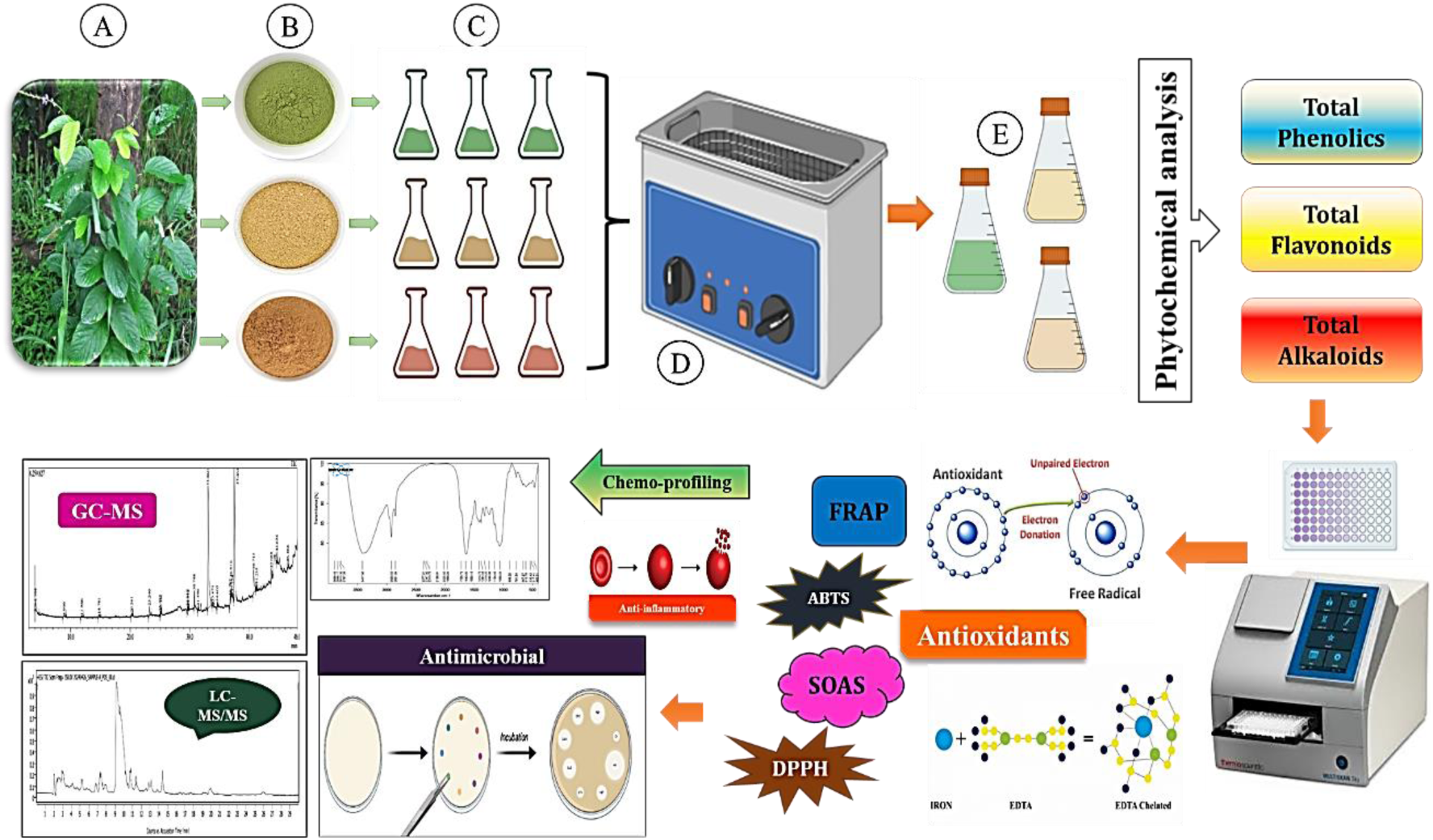
Graphical representation of Extraction, Phytochemical analysis, *in vitro* bioactivity profiling, and chemo profiling of *Desmodium gangeticum* (L.) DC. A: Leaves, stems and Roots of *D. gangeticum*; B: Fresh and Dry Powder; C: Extraction using three solvents Methanol, Ethanol, and Distilled Water; D: Extraction using Sonication; E: Prepared extracts of *D. gangeticum*.

### 2.4 Biochemical studies

#### 2.4.1 Quantification of Total Phenolic Content (TPC)

The total phenolic contents were determined by using the method described by Wolfe et al., (2003) with slight modifications. The reaction mixture had 25μL of extracts, 25μL Folin-Ciocalteu reagent, and 250μL ml of saturated Na_2_CO_3_ solution. The reaction mixture was allowed to stand at room temperature for 90 minutes. By using a Multiskan sky 96 well plate reader spectrophotometer, Thermo Scientific. Optical density was measured at 760 nm. The samples (in triplicates) were prepared for each analysis and an average of three absorbance readings were recorded. Calibration curve for standard phenolic compound Catechol was obtained by using a concentration of 20-100 µg/ml (r^2^= 0.998). Results were expressed as milligram Catechol equivalents (CAE) per gram fresh or dry weight (mg CAE/g FW/DW).

#### 2.4.2 Quantification of Total Flavonoid Content (TFC)

The total flavonoid contents of extracts were determined using the method reported by Luximon-Ramma et al., (2002). The reaction mixture contained 150μL of extract and 150μL of 2% methanolic AlCl_3_. The mixture was incubated for 10 minutes at room temperature, and the optical density (OD) was measured at 368 nm. For the experiment, the respective extraction solvent was used as a blank. A standard calibration curve for Rutin (20-100 µg/ml (r^2^= 0.946) was obtained. Results were expressed as milligrams of Rutin equivalent per gram of fresh or dry weight (mg RE/g FW/DW).

#### 2.4.3 Quantification of Total alkaloid Content (TAC)

Total alkaloid content (TAC) was evaluated using the 1, 10-phenanthroline method described by Singh et al., (2004). The assay mixture contained 100μL plant extract, 100μL 0.05M FeCl_3_ in 0.05M HCL, and 100μL 0.05M 1,10-phenanthroline reagent, final volume of reaction mixture was adjusted to 1 mL by using distilled water. This mixture was incubated for 15 minutes in water bath maintained at 75±2° C, till orange colour appears. The reaction mixture excluding plant extract was used as a blank. The absorbance was recorded at 510 nm. The absorbance was compared to a standard curve of a standard alkaloid Colchicine (r^2^=0.929). The values were expressed as milligrams of Colchicine equivalent (CE) per gram of fresh or dry weight (mg CE/g FW/DW).

### 2.5 Antioxidant activity

#### 2.5.1 Ferric Reducing Antioxidant Power Assay (FRAP)

The ferric ion reducing capacity was determined using the assay described by Pulido et al., (2000). To 10μl plant extract, 290μL of FRAP reagent [300 mM sodium acetate buffer at pH 3.6, 10 mM TPTZ solution and 20 mM FeCl_3_.6 H_2_O solution (10:1:1)] was added. The reaction mixture was incubated at 37^°^ C for 15 mins. The absorbance was measured at 595 nm. A calibration curve was prepared, using an aqueous solution of Ascorbic acid (20-100 µg/ml). The value of FRAP was expressed as micromolar of ascorbic acid equivalents per gram of plant sample (mM FRAP/g FW/DW).

#### 2.5.2 Phosphomolybdenum Reducing Power assay (PMo)

Total antioxidant capacity was assessed by Phosphomolybdenum method described by Prieto et al., (1999). The reaction mixture contains plant extract 25μL and 250μL of reagent solution (0.6 M sulphuric acid, 28 mM sodium phosphate and 4 mM ammonium molybdate). The reaction solution was incubated at 95° C for 90 min. After cooling to room temperature, the absorbance of solution was recorded at 695 nm. respective extraction solvent in the place of extract. was used as the blank. The PMo-reducing activity was calculated by preparing calibration curve using an aqueous solution of Ascorbic acid (20-100 µg/ml). The value of PMo was expressed as micromolar of ascorbic acid equivalents per gram of plant sample (mM PMo reduction/g FW/DW).

#### 2.5.3 Ferrous Ion Chelating Activity (FICA)

Ferrous ion chelating activity was determined using method of Dinis et al., (1994) with slight modification. Plant extract (100μL) mixed with 5μL of 2mM FeSO_4_, 185μL of distilled water was mixed into it. To start the reaction, 10μL of 5mM Ferrozine was added in that reaction. The mixture was incubated for 10 minutes at room temperature and absorbance was measured at 562 nm. Na_2_EDTA was used as positive control. Percent inhibition of Fe^+2^ ions of plant extracts were calculated using the following formula:

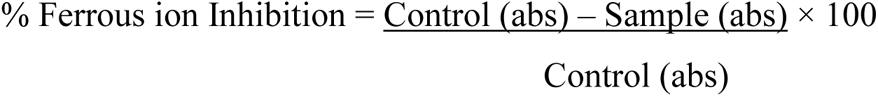

#### 2.5.4 Superoxide Anion Scavenging Assay (SOAS)

Superoxide anion scavenging activity assay performed by using method of Tiwari et al., 2017. The reaction mixture was prepared by adding 20μL of (NBT) Nitroblue tetrazolium (1 mg of NBT in 1 ml DMSO), 60μL of plant extract, and 200μL of alkaline (DMSO) dimethyl sulfoxide (1 ml of alkaline DMSO containing 0.1 ml of 5 mM NaOH and 0.9 ml of DMSO). The absorbance was recorded at 560 nm. DMSO solution served as blank. The SOAS activity was calculated by using the formula:

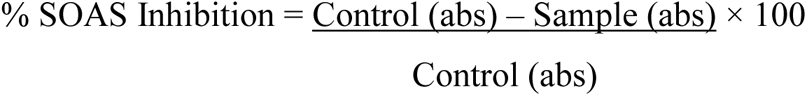

#### 2.5.5 ABTS (2, 2’-azino-bis (3-ethylbenzthiazoline-6-sulphonic acid) scavenging activity

ABTS activity was assayed by using a method studied by Re et al., 1999. Plant extract (10 µl, 1%) was mixed with 290 µl of ABTS reagent. An aqueous solution of ABTS (7mM) was mixed with a 2.45mM aqueous solution of potassium persulfate in a 1:1 proportion. This mixture was incubated in the dark for 12-16 hrs. Working solution of ABTS was prepared by diluting the reagent to give an absorbance 0.7±0.02 at 734 nm. The reaction mixture was incubated at room temperature for 10 mins and scavenging activity was measured by checking absorbance at 734 nm. Ascorbic acid was used as standard. ABTS served as a control. The percentage of inhibition or scavenging was calculated by,

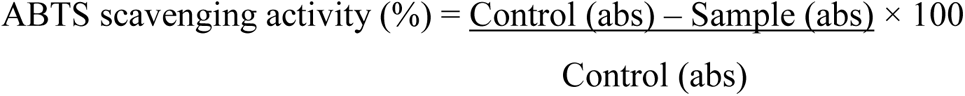

#### 2.5.6 DPPH (2, 2-diphenyl-1-picrylhydrazyl) radical scavenging assay

The free radical scavenging activity of plant extract was measured as per method of Aquino et al., (2001). Plant extract (20μL) was mixed with 200μL of 25mM methanolic DPPH solution. The reaction mixture was incubated in the dark at room temperature for 30 min. The absorbance was measured at 517 nm. DPPH served as a control. Results were expressed as percentage inhibition of the DPPH radical and was calculated using the following formula:

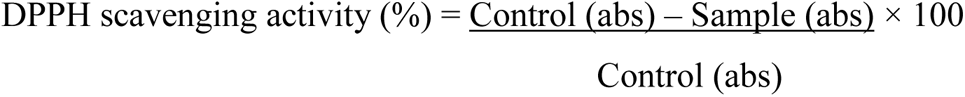

### 2.6 Anti-inflammatory activity

#### 2.6.1. HRBC membrane stabilization activity

##### 2.6.1.1 Preparation of Red Blood cells suspension (RBCs)

In their study on anti-inflammatory activity, Mane et al., (2022) detailed the process of preparing a suspension of red blood cells (RBCs). Fresh human blood (5 mL) was obtained from a participant who refrained from NSAIDs for a minimum of two weeks before the experiment. EDTA was employed as an anticoagulant during blood collection to prevent clotting. The sample was centrifuged at 3000 rpm for ten minutes, followed by two washes with an equivalent volume of normal saline (0.9% NaCl). Finally, a 10% v/v suspension of the blood cells was meticulously prepared using an isotonic buffer solution (10 mM sodium phosphate buffer, pH 7.4, containing 154 mM NaCl).

##### 2.6.1.2. Hypotonic solution-induced hemolysis

Mane et al., (2022) investigated the impact of hypotonic solution-induced hemolysis on the membrane-stabilizing effect of plant extracts. Initially, 125 μL of either the sample or a standard acetylsalicylic acid (ASA) as a positive control were added to a 1.5 ml centrifuge tube containing 1 ml of hypotonic solution (25 mM NaCl in 10mM sodium phosphate buffer, pH 7.4). Following this, 125 μL of a red blood cell (RBC) suspension was mixed into the reaction mixture and left at room temperature for 10 minutes. A control was set up with 1.125 ml of hypotonic solution and 125 μL of RBC suspension. Subsequently, the samples were centrifuged for five minutes at 2500 rpm, and the absorbance of the supernatant was measured at 540 nm. The percentage inhibition of hemolysis or membrane stabilization was calculated using the formula:

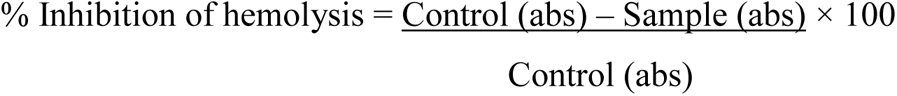

### 2.7 Antibacterial Activity

Mutyala and Owk (2016) studied antibacterial activity using the Agar disc diffusion method. The microbial strains employed in this study were gram-positive bacteria *Staphylococcus aureus* (NCIM 2654) and gram-negative bacteria *Escherichia coli* (NCIM 2832).

#### 2.7.1 Media preparation: Mueller Hinton Agar Medium

To formulate the Mueller Hinton Agar medium, 38 grams of Mueller Hinton Agar (Himedia) powder were accurately measured and dissolved in 1000 millilitres of distilled water. The resulting solution was sterilized by autoclaving at 15 pounds of pressure and 121°C for 15 minutes. After autoclaving, the medium was allowed to cool, mixed thoroughly, and aseptically poured into sterile petri plates.

#### 2.7.2. Preparation of Bacterial Suspension

The inoculum was prepared by culturing bacteria in 25 mL of Luria Bertani (LB) broth and incubating it overnight at 37°C until the OD_600_ reached an absorbance equivalent to 1.0, as determined by the growth curve of the microorganisms. The antimicrobial activity of methanol, ethanol, and aqueous extracts from the leaves, stems, and roots (both fresh and dried) of *D. gangeticum* was assessed using the agar disc diffusion method. The plant extracts were diluted in their respective solvents in a serial two-fold dilution, ranging from 6.25 milligrams per millilitre to 100 milligrams per millilitre. Sterilized blank discs were immersed in the respective plant samples for 4 hours.

Under aseptic conditions, bacteria were inoculated into LB broth and then incubated at 37°C for 18 hours. Following the incubation period, 100 microliters of the test bacterial culture was pipetted onto Mueller Hinton Agar plates using a micropipette and spread evenly with a spreader. Five discs, each dipped in the respective solvents, were placed on each plate. Positive controls were established by including sterilized discs with an equivalent quantity of the antibiotic streptomycin. Respective extraction solvent was used as a blank. The petri dishes were subsequently incubated at 37°C for 24 hours. The antimicrobial activity was assessed by measuring the diameter of inhibition zones around the discs, recorded in millimetres.

### 2.8. Fourier-Transform Infrared spectrometry (FTIR)

FTIR spectra were acquired using an Alpha Bruker FT-IR spectrometer. A small amount of each sample powder was placed directly on KBr pellets under constant pressure. Infrared absorbance data were collected across the wave number range of 4000 cm⁻¹ to 400 cm⁻¹ to analyze the frequencies of functional groups in the samples. Spectral analysis focused on identifying peak values within the infrared radiation region for all samples.

### 2.9. Gas Chromatography and Mass Spectrometry (GC-MS/MS)

GC–MS/MS analyses of leaf and root extracts of methanol were carried out using the TQ 8050 plus with HS-20 (Shimadzu, Japan) equipped with a SHIMADZU SH-Rxi-5Sil MS column (30 m in length × 0.25mmID × 0.25 μm df). The carrier gas was pure helium gas (99.99%) at a consistent rate of 1 mL/min. Fragmentation ranging from 45 to 500 m/z was used for GC–MS/MS spectrum identification. A 1 μl injection volume was utilized, and a constant injector temperature of 250 °C was maintained. The temperature of the column oven was initially set at 50 °C and then increased to 260 °C. This final temperature was kept constant. By comparing the test samples mass, peak area, peak height, and retention time (min) with the authentic spectral compounds databases kept in the National Institute of Standards and Technology (NIST) library, the phytochemicals present in them were determined.

### 2.10. Metabolite profiling of the plant crude extract by UPLC-QTOF-MS

Metabolite profiling of the crude extract was done using Agilent Q-ToF G6540B connected to Agilent 1260 Infinity II HPLC in positive mode (ESI+). Filtering distilled water extracts of dried leaf and root were analysed 0.25 mm polyvinylidene fluoride (PVDF) membrane syringe filters into a 2 mL vial. A sample injection volume of 5 µl was used for chromatographic separation of analytes through an Agilent Eclipse XDB-C18, 3X150 mm, 3.5-micron column. The analytical run was set at 30 mins, and the MS Scan Range at *m/z* 100 to 1700. The mobile phase comprised two solvents: solvent (A) deionized water containing 0.1 % formic acid and solvent (B) acetonitrile containing 0.1 % formic acid, following the chromatographic gradient as shown in Table 1.

**Table 1:**
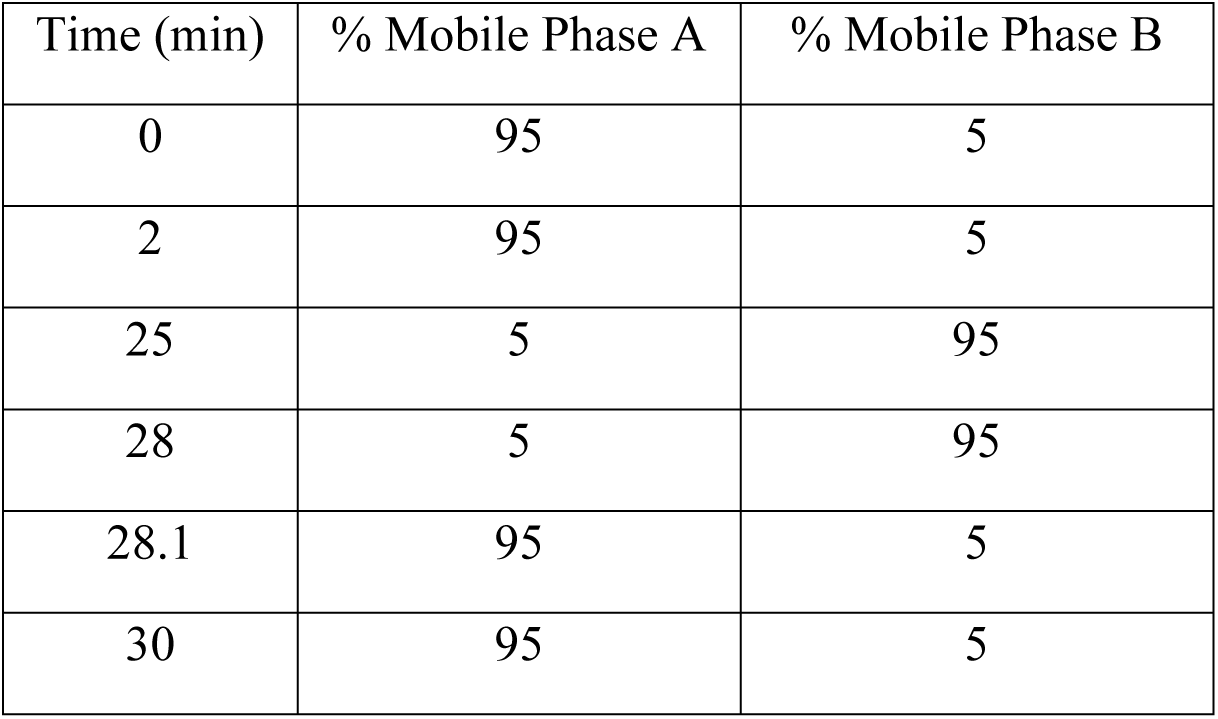
Chromatographic gradient (Flow profile) of UPLC-QTOF-MS for *D. gangeticum*.

The elution was carried out at room temperature at a flow rate of 0.3 ml/min. The flow profile of the mobile phase is mentioned in Table 1. The run was carried out at 300°C Nebulizer Gas Temperature, 350°C Sheath Gas Temp, 8 l/min Drying Gas, 35 psig Nebulizer Gas, 11 l/min Sheath Gas Flow, 3500V Capillary voltage, 1000V Nozzle Voltage Ion Source Parameters.

### 2.11. Statistical Analysis

The average values of triplicate experiments were used for statistical analysis, and the data are expressed as mean ± standard error. The significant cut-off point was set at P<0.05. The data was analysed using GraphPad Prism 5 Software, San Diego, California, USA. The primary results were also computed using Microsoft Excel formulas.

## 3. Results and Discussion

### 3.1 Biochemical Analysis

#### 3.1.1 Total Phenolic Content

Phenolics are a group of secondary metabolites found in a diverse range of plants, known for their antioxidants (Pandey and Rizvi, 2009; Scalbert et al., 2005), antimicrobial, and other bioactive properties. In the present investigation, the total phenolic content in *D. gangeticum* was assessed using various solvents (methanol, ethanol, and distilled water) as shown in Fig. 2. The highest phenolic content was observed in the dried leaf powder extracted with distilled water with 8.02 ± 0.01 mg CAE/g DW. This was followed by the methanol extract of dried roots (2.48 ± 0.005 mg CAE/g DW) and the ethanol extract of dried leaves (1.16 ± 0.003 mg CAE/g DW). These results suggest that extracts obtained with distilled water contain higher levels of phenolics than those obtained with ethanol and methanol across all plant parts of *D. gangeticum*. Phenols are utilized in various applications, including medications, oral analgesics, antiseptics for surgical tools, skincare products, and embalming processes. The diverse array of active chemicals in *D. gangeticum* plays a crucial role in its potential anticancer properties (Srivastava et al., 2013; 2015); however, this assertion would benefit from explicit references to specific studies and the underlying mechanisms of action that support these claims. Notably, three phenolic compounds— methyl salicylate β-D-glucopyranoside, syringaresinol-4’-O-β-D-glucopyranoside, and leonuriside A —were isolated from a water-soluble extract of *D. gangeticum* leaves (Dat et al., 2015).

**Fig. 2.**
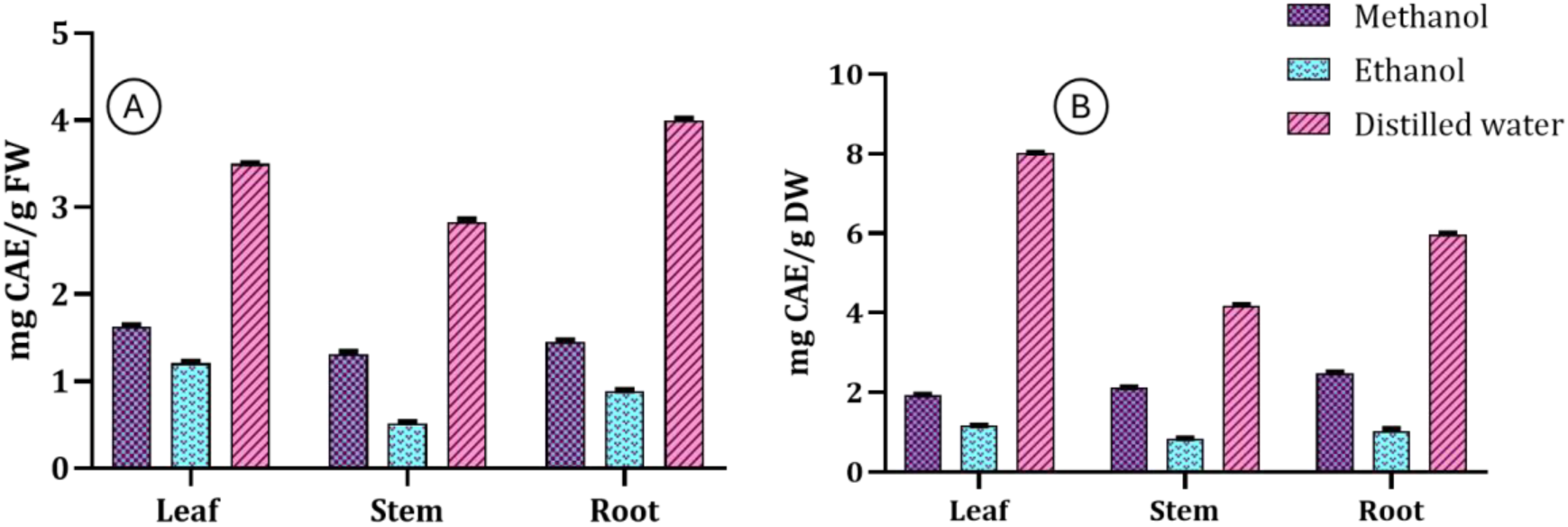
Total phenolic contents in *D. gangeticum* in different solvents (methanol, ethanol and aqueous) A) from Fresh plant (leaf, stem and root), B) from Dried plant (leaf, stem and root). The results were reported as mean values ± SE (P < 0.05).

#### 3.1.2 Total Flavonoid Content

Flavonoids are secondary metabolites known for their antioxidant potential and significant roles in food, cosmetics, medicine, and biological processes (S. Chen et al., 2023). They are well-documented for their antioxidant activity, effectively scavenging free radicals and protecting cells and tissues from oxidative damage, thus making them valuable in nutritional and health applications. The total flavonoid contents are shown in Fig. 3. The highest flavonoid content was observed in dried leaves extracted with distilled water (45.40 ± 0.02 mg RE/g DW), while the lowest was found in the ethanolic extract of fresh stems (3.57 ± 0.01 mg RE/g FW). The methanolic extract of dried leaves recorded 21.15 ± 0.02 mg RE/g DW, and the ethanolic extract of dried leaves showed 20.46 ± 0.04 mg RE/g DW. Among the plant parts, dried leaf extracts demonstrated a higher flavonoid content across all solvents studied. Distilled water proved to be the most efficient solvent for extracting flavonoids from leaves, stems, and roots compared to methanol and ethanol. Our findings indicate higher flavonoid content than previously reported by Venkatachalam and Muthukrishnan (2012), who found 10.5 ± 0.5 mg CAE/mL in ethanol extracts of *D. gangeticum*. Tsai et al., (2011) reported that the whole plant extract of *D. gangeticum* in 70% ethanol contained 127.49 ± 2.73 µg RE/mg DW of flavonoids. Flavonoid content varied significantly depending on the solvent and plant parts used for extraction, with dried plant parts showing higher flavonoid content in methanol and ethanol compared to distilled water. The results indicate that leaves generally have higher flavonoid content than stems and roots. Thus, the choice of solvents and plant parts significantly impacts flavonoid extraction efficiency.

**Fig. 3.**
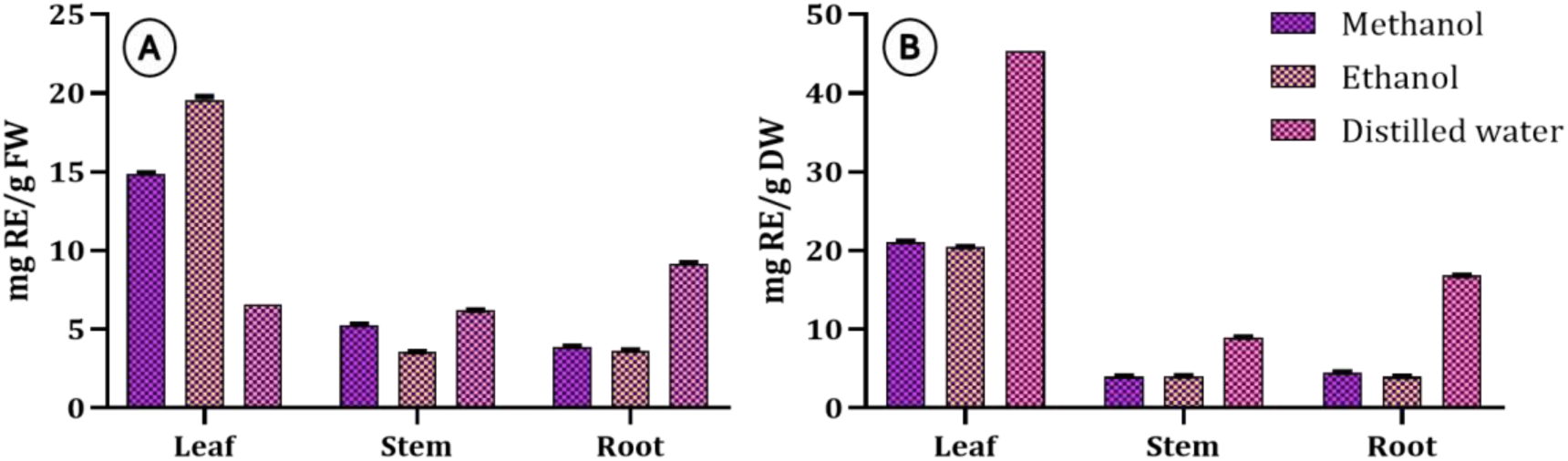
Total flavonoid contents in *D. gangeticum* in different solvents (methanol, ethanol and aqueous) A) from Fresh plant (leaf, stem and root), B) from Dried plant (leaf, stem and root). The results were reported as mean values ± SE (P < 0.05).

#### 3.1.3 Total Alkaloid Content

Alkaloids are nitrogen-containing compounds renowned for their diverse pharmacological activities. *D. gangeticum*, known for its medicinal properties in traditional folk medicine, contains a notable concentration of alkaloids. The highest alkaloid content was found in methanol extracts across both fresh and dried samples, suggesting that methanol may be more efficient in extracting alkaloids from *D. gangeticum* compared to ethanol and distilled water. The study also reveals considerable variation in alkaloid distribution among different plant parts (leaf, stem, root), with leaves being more abundant in alkaloids compared to stems and roots. The dried leaves extracted in methanol showed the highest alkaloid content, 25.36 ± 0.13 mg CE/g DW, while the lowest was found in the ethanolic extract of fresh stems (4.85 ± 0.04 mg CE/g FW) (Fig. 4).

**Fig. 4.**
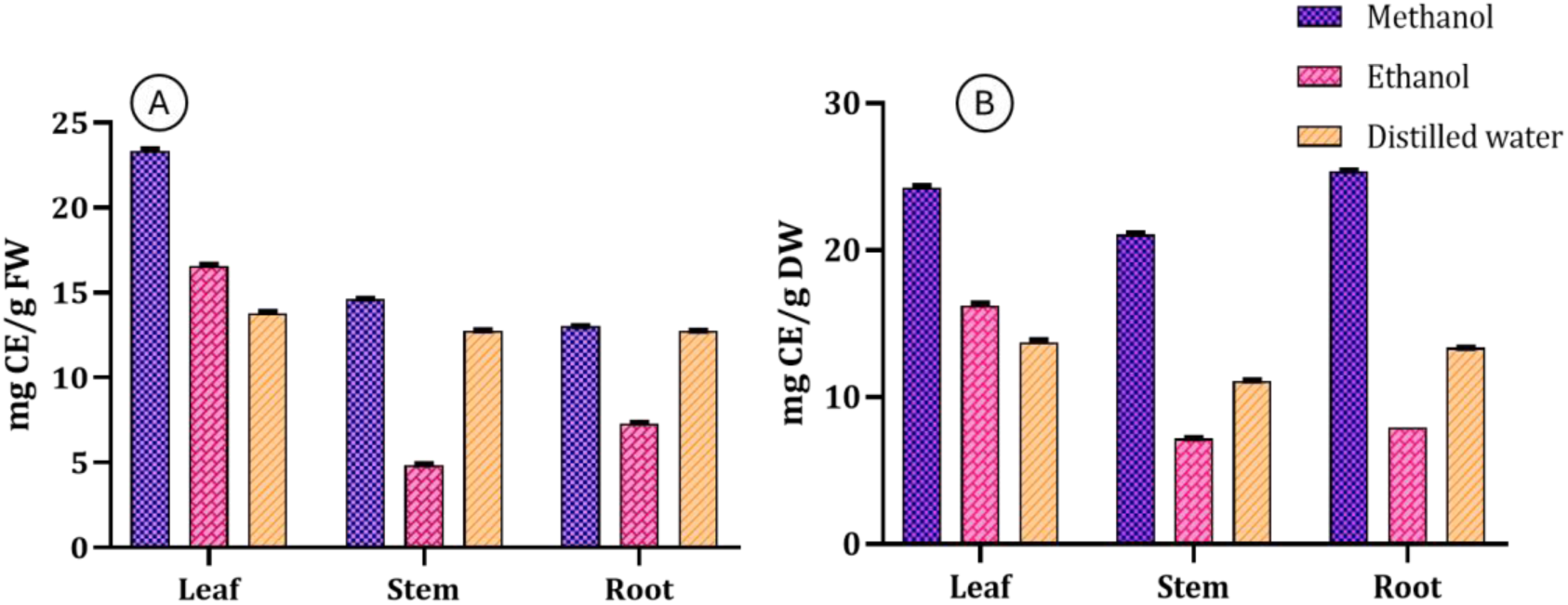
Total alkaloid contents in *D. gangeticum* in different solvents (methanol, ethanol and aqueous) A) from Fresh plant (leaf, stem and root), B) from Dried plant (leaf, stem and root). The results were reported as mean values ± SE (P < 0.05).

Alkaloids are complex nitrogenous substances with high molecular weights present in microbes, plants, and animals (Masci et al., 2019). A study by Mahajan et al., (2015) demonstrated the anti-amnesic efficacy of an alkaloidal fraction of *D. gangeticum* highlighting the therapeutic potential of its alkaloid profile. This finding is significant as it underscores the pharmacological versatility of alkaloids present in *D. gangeticum*, which are not only associated with neuro but may also contribute to a range of other pharmacological activities, including antianxiety and analgesic properties. By linking the specific effects of alkaloids to cognitive health this study emphasizes the importance of further exploring the diverse roles of alkaloids in traditional medicines and their potential application in modern therapeutics. Subsequent investigations have demonstrated its antianxiety, anticonvulsant, antidepressant, locomotor, and hypnotic activities. Additionally, Changdar et al., (2019) inter-related the neuroprotective activity of *D. gangeticum* to its alkaloids. Previous studies have highlighted the presence of alkaloids in *D. gangeticum* and their diverse pharmacological characteristics, including analgesic, anti-inflammatory, and neuroprotective properties.

#### 3.2.1 Ferric Reducing Antioxidant Power (FRAP) Activity

The Ferric Reducing Antioxidant Power (FRAP) assay measures the antioxidant capacity of substances based on their ability to reduce ferric ions (Fe³⁺) to ferrous ions (Fe²⁺) through an electron transfer mechanism. Antioxidants donate electrons to ferric ions, leading to the formation of ferrous ions which can be quantified spectrophotometrically (Wojtunik-Kulesza, 2020). The FRAP assay was used to assess the antioxidant potential of *D. gangeticum* extracts from various plant parts and solvents. Results revealed significant variation in antioxidant potential different extracts (Fig. 5). Distilled water extracts consistently showed higher FRAP values compared to methanol and ethanol extracts, indicating a higher concentration of water-soluble antioxidants. Specifically, the highest FRAP activity was found in the distilled water extract of dried leaf powder, with a value of 10.81 ± 0.03 mM FRAP/g DW. This suggests that *D. gangeticum* possesses substantial antioxidant properties, particularly in combating oxidative stress-related disorders. The presence of flavonoids and polyphenols in these extracts likely contributes to their strong antioxidant activity by scavenging free radicals.

**Fig. 5.**
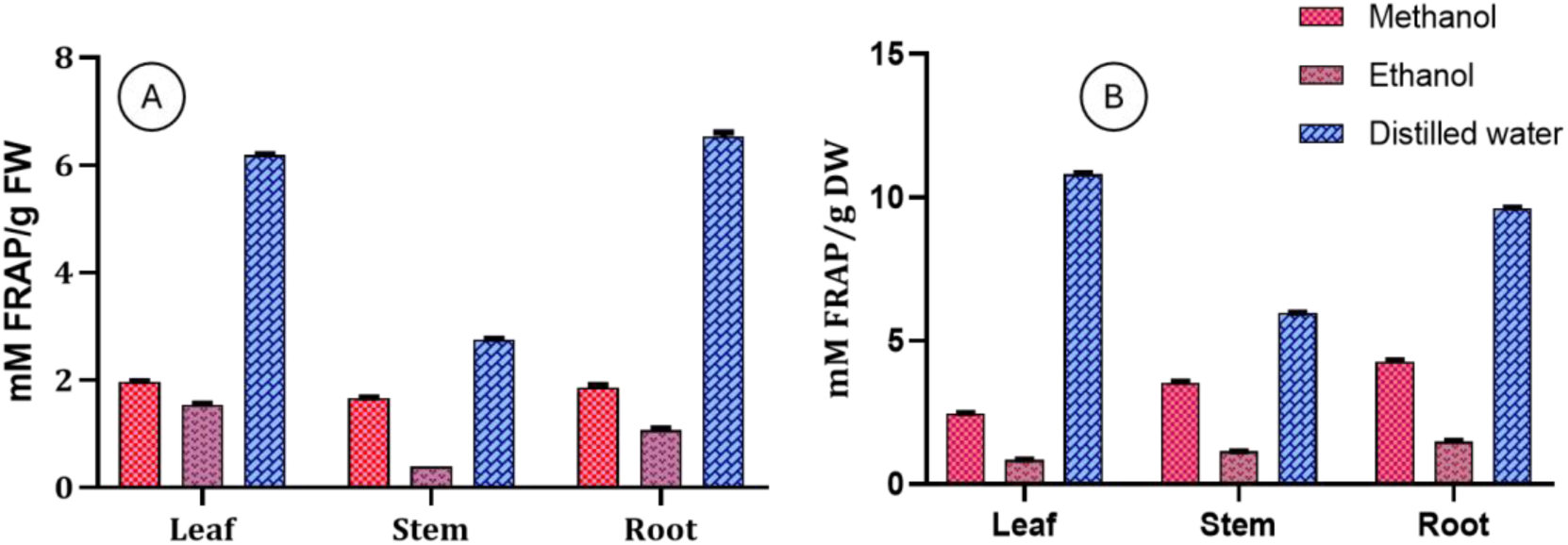
FRAP activity in *D. gangeticum* in different solvents (methanol, ethanol and aqueous) A) from Fresh plant (leaf, stem and root), B) from Dried plant (leaf, stem and root). The results were reported as mean values ± SE (P < 0.05).

#### 3.2.2 Phosphomolybdenum Reducing Power Assay (PMo)

The Phosphomolybdenum (PMo) assay evaluates the reducing capacity of antioxidants by measuring the reduction of Mo (VI) to Mo (V) and the formation of a green phosphate-Mo (V) complex at acidic pH (Ravishankar et al., 2014). In this study, the reducing capacity varied across plant parts and solvents. The highest PMo activity was observed in the distilled water extract of dried leaves, recording 6.26 ± 0.005 mM PMo reduction/g DW. This was followed by methanol (5.30 ± 0.004 mM PMo reduction/g DW) and ethanol (5.01 ± 0.004 mM PMo reduction/g DW) extracts of dried leaves (Fig. 6). These results indicate that water-soluble compounds, such as phenolics and flavonoids, which are known for their electron-donating abilities, contribute to the high reducing power. The leaf extracts, especially from dried material, exhibited greater reduction potential.

**Fig. 6.**
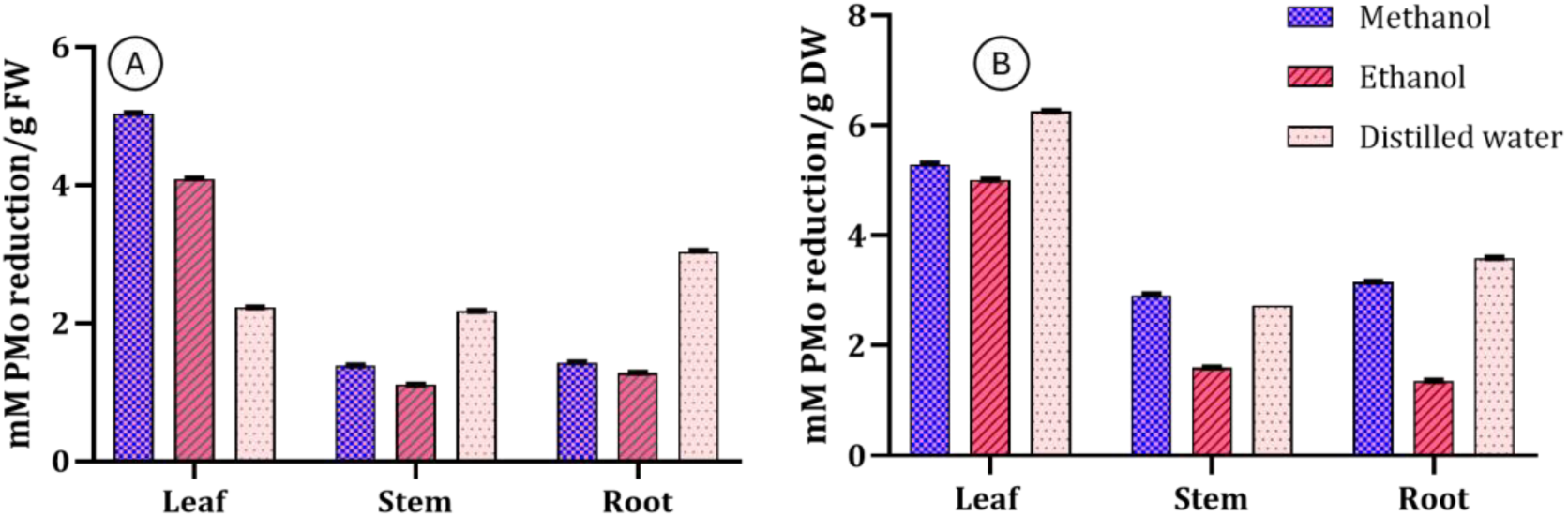
PMo activity in *D. gangeticum* in different solvents (methanol, ethanol and aqueous) A) from Fresh plant (leaf, stem and root), B) from Dried plant (leaf, stem and root). The results were reported as mean values ± SE (P < 0.05).

#### 3.2.3 Ferrous Ion Chelating Activity

Ferrozine and Fe²⁺ form a red complex, which can be quantitatively measured. Antioxidants inhibit this process by chelating ferrous ions, thus reducing the color intensity of the ferrozine-Fe²⁺ complex. This inhibition reflects the chelating activity of the antioxidants (Soler-Rivas et al., 2000). The highest ferrous ion chelating activity, with an inhibition of 67.44 ± 0.02%, was observed in the distilled water extract of fresh stems (Fig. 7). The chelating capacity of the ethanolic extract of *D. gangeticum* was previously assessed by Venkatachalam and Muthukrishnan (2012), who reported a chelating ability ranging from 21.12% to 73.99% depending on the concentration. EDTA exhibited strong chelating activity. Our findings support the notion that the bioactive constituents of *D. gangeticum* contribute significantly to its metal binding properties and antioxidant capacity.

**Fig. 7.**
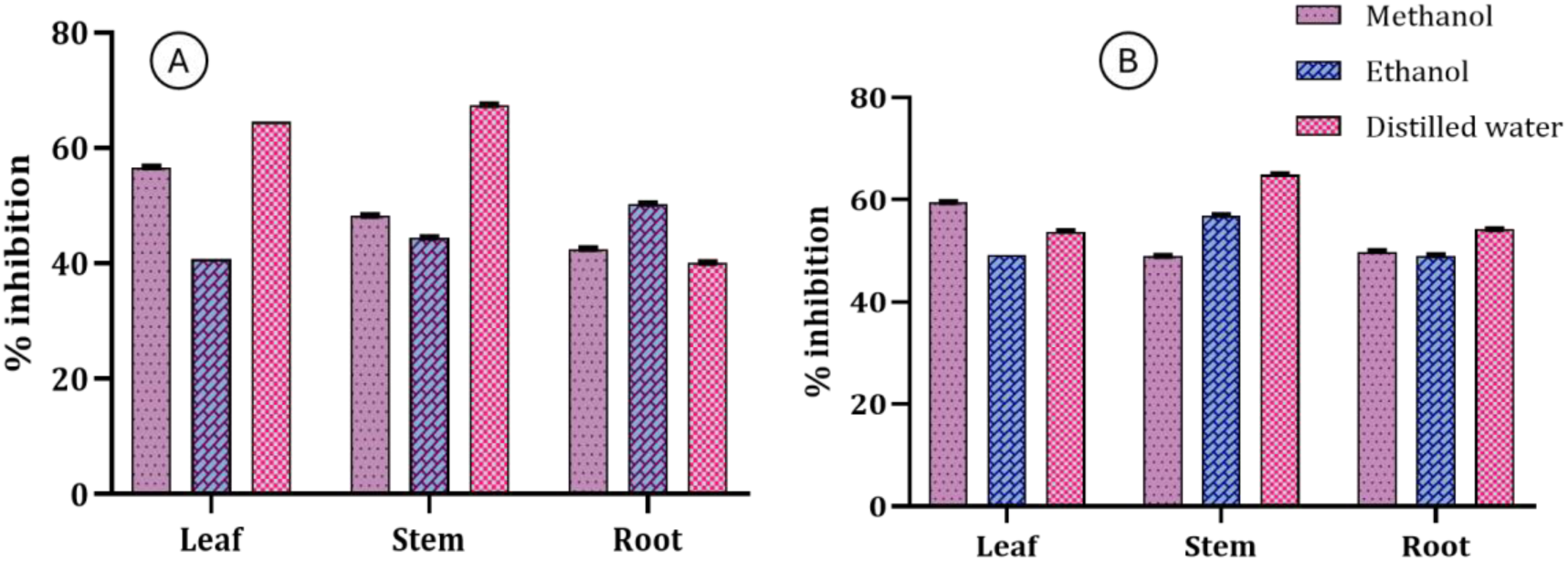
FICA activity in *D. gangeticum* in different solvents (methanol, ethanol and aqueous) A) from Fresh plant (leaf, stem and root), B) from Dried plant (leaf, stem and root). The results were reported as mean values ± SE (P < 0.05).

#### 3.2.4 Superoxide Anion Scavenging Activity (SOAS)

Superoxide anion scavenging activity is crucial for mitigating oxidative stress and maintaining cellular integrity. This activity reflects the capacity of substances to neutralize superoxide radicals (O₂⁻), which are reactive oxygen species (ROS) involved in various pathological conditions (Andrés et al., 2023). The methanol, ethanol, and aqueous extracts of *D. gangeticum* demonstrated significant inhibition of superoxide generation. The highest scavenging activity was observed in the ethanolic extract of fresh stems, with 90.85% inhibition (Fig. 8). All three fresh plant parts showed notable scavenging activity in methanol, ethanol, and distilled water. Kurian et al., (2010b) reported a 92.31 ± 2.63% inhibition of superoxide radicals by an ethyl acetate extract of the entire plant, which aligns with our results. The findings suggest that *D. gangeticum* is a promising medicinal plant for scavenging superoxide anions.

**Fig. 8.**
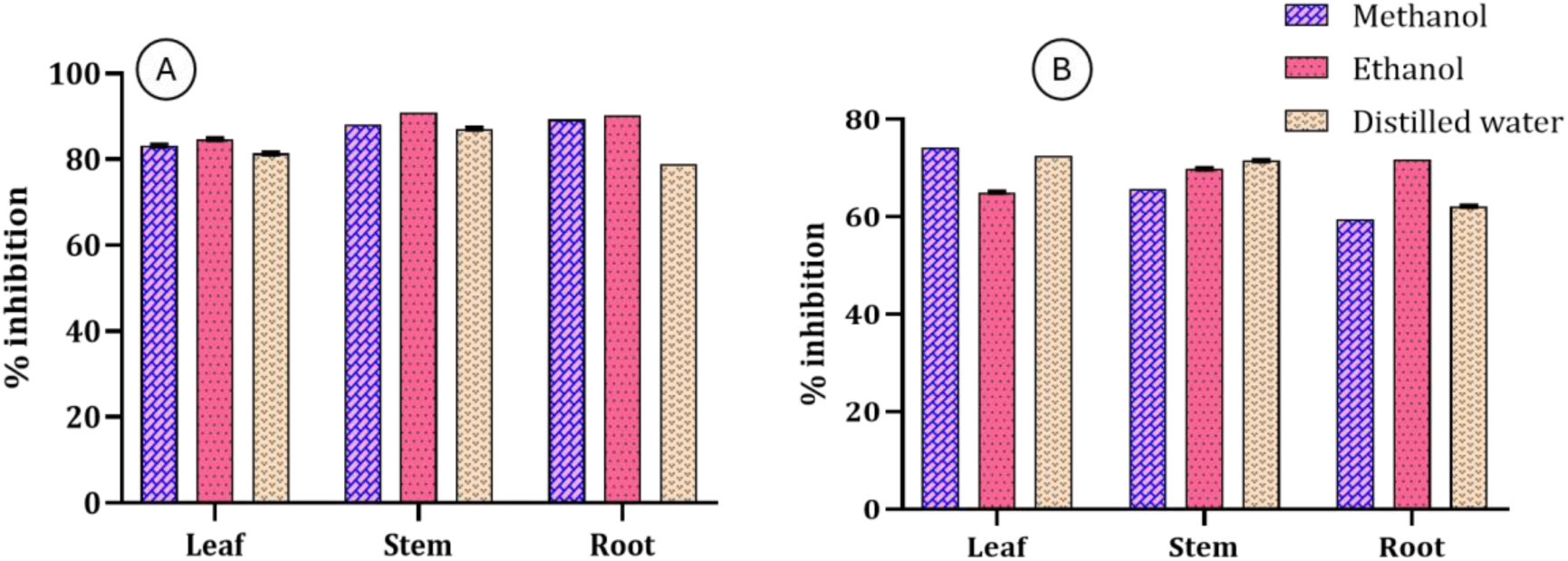
SOAS activity in *D. gangeticum* in different solvents (methanol, ethanol and aqueous) A) from Fresh plant (leaf, stem and root), B) from Dried plant (leaf, stem and root). The results were reported as mean values ± SE (P < 0.05).

#### 3.2.5 ABTS Radical Scavenging Activity

The ABTS assay measures the ability to reduce ABTS⁺ free radicals, resulting in a decrease in color intensity. The extent of discoloration indicates antioxidant capacity (Sabbione et al., 2016). The highest ABTS radical scavenging activity was observed in the distilled water extract of dried leaves, showing 90.44 ± 0.005% inhibition (Fig. 9). This suggests the presence of water-soluble compounds like phenols and flavonoids, which effectively scavenge free radicals. Mathew and Abraham (2004) highlighted the role of antioxidants in scavenging proton radicals, with the protonated ABTS radical exhibiting a maximum absorbance at 734 nm, which decreases upon scavenging. Our results are consistent with their findings, showing that the leaf ethanolic extract of *D. gangeticum* is a potent scavenger of ABTS radicals.

**Fig. 9.**
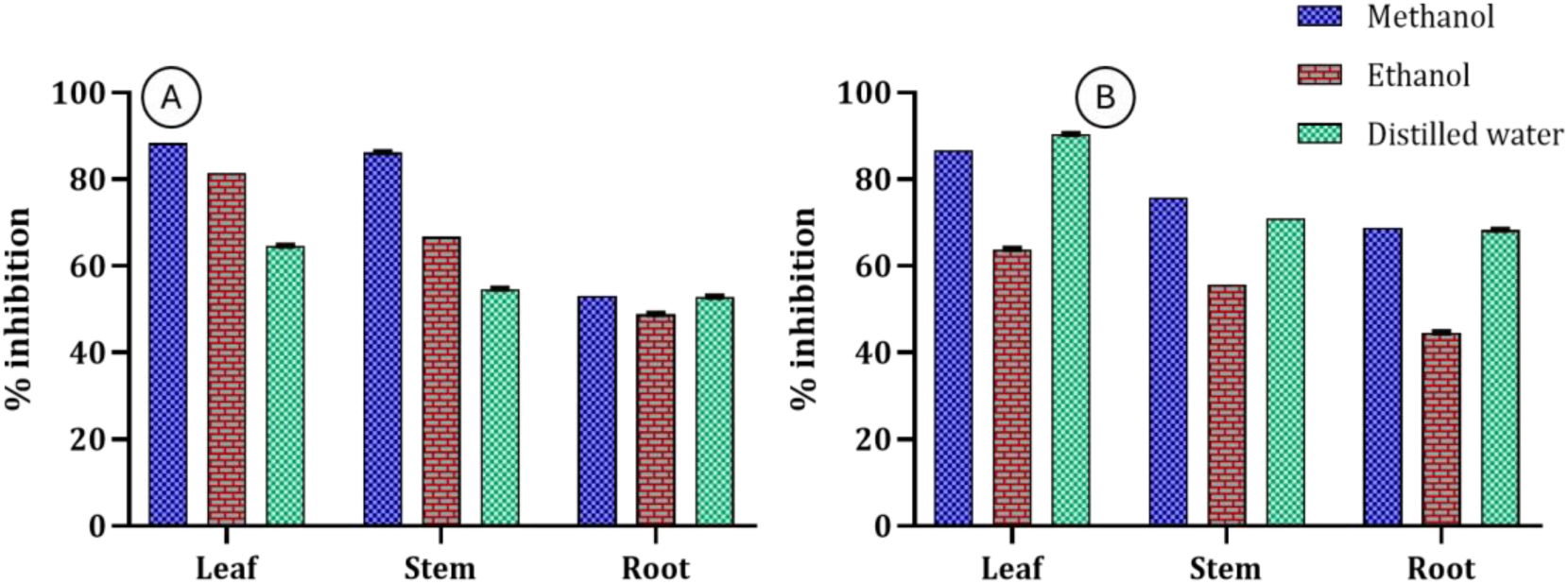
ABTS activity in *D. gangeticum* in different solvents (methanol, ethanol and aqueous) A) from Fresh plant (leaf, stem and root), B) from Dried plant (leaf, stem and root). The results were reported as mean values ± SE (P < 0.05).

#### 3.2.6 DPPH Radical Scavenging Assay

The DPPH scavenging assay uses the stable free radical DPPH, which has a deep violet color. Antioxidants in plant extracts neutralize DPPH by donating electrons or hydrogen, leading to a color change from violet to yellow or pale yellow. The degree of discoloration reflects the scavenging capacity (Venkatachalam and Muthukrishnan, 2012). The methanolic extract of dried roots showed the highest DPPH scavenging activity, with 65.18 ± 0.10% inhibition (Fig. 10). Methanol was found to be more effective in scavenging DPPH compared to distilled water and ethanol. Although the root exhibited the highest activity, Kurian et al., (2010b) reported 89.25 ± 2.11% DPPH radical scavenging activity for the entire plant extracted in ethyl acetate.

**Fig. 10.**
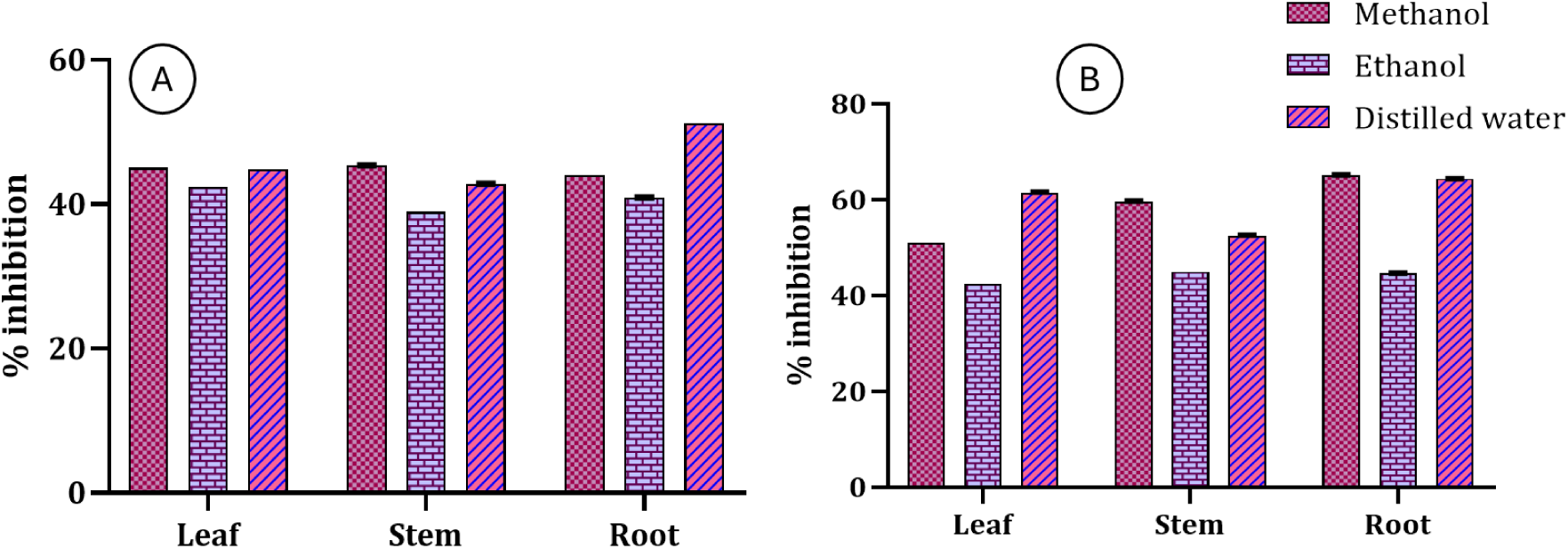
DPPH activity in *D. gangeticum* in different solvents (methanol, ethanol and aqueous) A) from Fresh plant (leaf, stem and root), B) from Dried plant (leaf, stem and root). The results were reported as mean values ± SE (P < 0.05).

The results demonstrate that *D. gangeticum* possesses significant antioxidant activity, which may underpin its diverse biological and therapeutic properties. The data indicates a positive relationship between the bioactive compounds (particularly phenolics and flavonoids) and the antioxidant activities of *D. gangeticum*. Higher concentrations of these compounds in specific extracts (e.g., distilled water extracts of dried leaves) are associated with stronger antioxidant activity, supporting the idea that these compounds contribute significantly to the plant’s overall antioxidant profile. The antioxidant activity is likely due to the synergistic effects of polyphenolic compounds, including chlorogenic and caffeic acids. Previous studies on *D. gangeticum* have also highlighted its potent antioxidant properties (Kumar et al., 2014; Pramanik et al., 2016; Govindarajan et al., 2003). Methanolic extracts have shown considerable antioxidant potential, but our study indicates that both aqueous and methanolic extracts are effective, likely due to the phytocompounds extracted in these solvents.

### 3.3 Anti-inflammatory Activity

Non-steroidal anti-inflammatory drugs (NSAIDs) protect cell membranes by stabilizing lysosomal membranes and preventing the release of lysosomal enzymes, thereby reducing cell damage and fluid leakage (Magdum et al., 2024; Mounnissamy et al., 2007). The anti-inflammatory activity of *D. gangeticum* was evaluated by assessing its ability to inhibit hypotonicity-induced hemolysis of human red blood cells (HRBCs), a model for lysosomal membrane stabilization. HRBC membranes, which share similarities with lysosomal membranes, were used to gauge the anti-inflammatory potential of the extracts.

In this study, the distilled water extract of fresh leaves exhibited the highest hemolysis inhibition, followed by the stem and root extracts, with inhibition percentages of 54.88 ± 0.002%, 54.77 ± 0.003%, and 53.68 ± 0.001%, respectively (Fig. 11). These results indicate that all parts of *D. gangeticum*, both fresh and dried, possess notable membrane-stabilizing properties. Previous research has shown that *D. gangeticum* extracts exhibit significant anti-inflammatory and analgesic effects in various animal models (Sagar et al., 2010; Rathi et al., 2004). This study provides novel insights into the membrane stabilization activity of *D. gangeticum* extracts across different solvents and plant parts, which has not been previously reported.

**Fig. 11.**
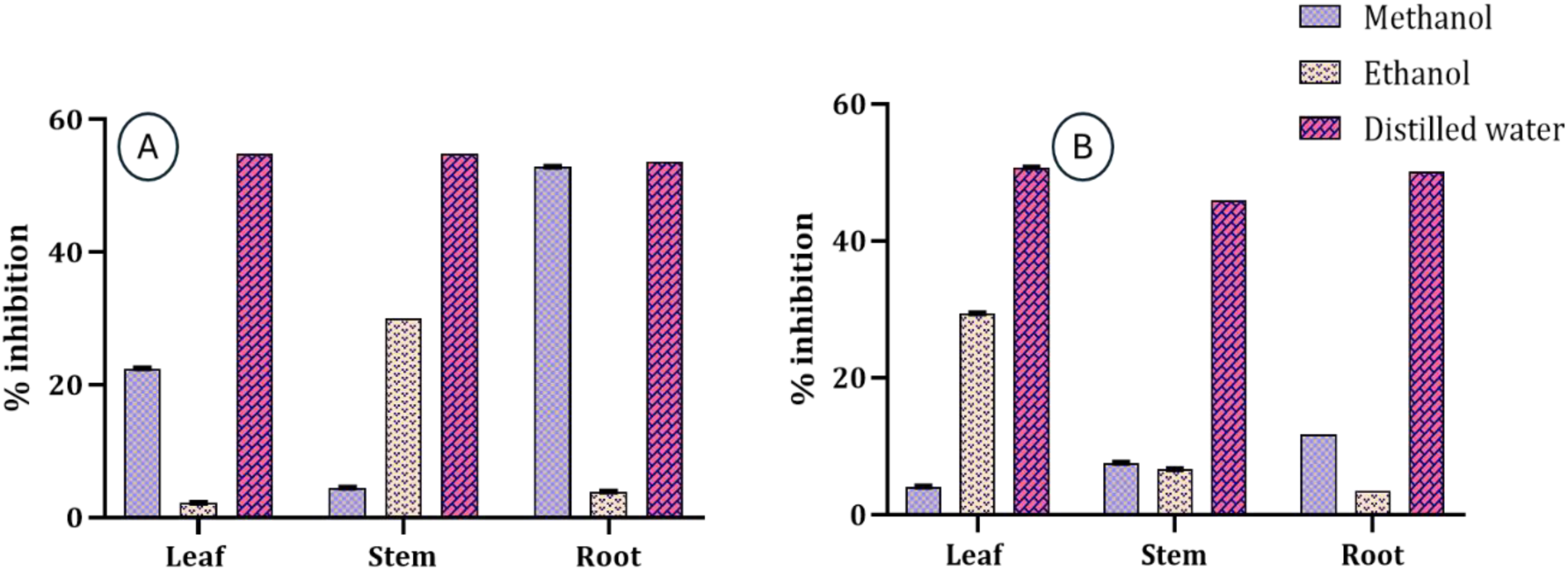
Anti-inflammatory activity in *D. gangeticum* in different solvents (methanol, ethanol and aqueous) A) from Fresh plant (leaf, stem and root), B) from Dried plant (leaf, stem and root). The results were reported as mean values ± SE (P < 0.05).

### 3.4 Antibacterial Activity

The antibacterial efficacy of *D. gangeticum* extracts was assessed using the disc diffusion method, with inhibition zones measured in millimeters (mm). The highest inhibition zone against the Gram-positive bacterium *Staphylococcus aureus* was observed with the distilled water extract of dried leaves at 100 mg/ml, showing a diameter of 11.5 ± 0.9 mm (Supplementary table 1). For the Gram-negative bacterium *Escherichia coli*, the fresh leaf extract in distilled water exhibited the greatest inhibition zone of 10 ± 0.01 mm (Supplementary table 2). Both distilled water and ethanol extracts showed antimicrobial activity, while methanol extracts demonstrated comparatively lower activity. Streptomycin, used as a reference antibiotic, showed the largest inhibition zone of 27 ± 0.1 mm at 100 mg/ml.

Krishnasamy et al., (2012) reported that the methanolic extract of *D. gangeticum* had a maximum zone of inhibition of 24 ± 2.3 mm against *Streptococcus mutans*, whereas aqueous extracts showed minimal activity against *Pseudomonas aeruginosa* (7 ± 0.08 mm). Hemlal and Subban (2012) noted that methanolic extracts of *D. gangeticum* effectively inhibited various pathogens, including *Bacillus cereus* and *Aeromonas hydrophila*, with inhibition zones of 12.0 mm and 13.0 mm, respectively. Mutyala and Owk (2016) found that methanol extracts were most effective against *Pseudomonas aeruginosa*, with comparable efficacy against *Bacillus subtilis* and *Escherichia coli*. Present findings validate previous studies but reveal some noteworthy discrepancies. Methanol extracts showed limited antibacterial activity against both tested bacteria. However, distilled water extracts of *D. gangeticum* exhibited significant antibacterial activity, comparable to that of Streptomycin. Given the rise in infections and antibiotic resistance (Austin et al., 1999), exploring natural antimicrobial sources is increasingly vital. Traditional medicinal systems like Unani and Ayurveda have long utilized plant-derived remedies for treating bacterial and fungal infections (Mutyala and Owk, 2016). This study emphasizes the potential of *D. gangeticum* as a source of effective natural antibiotics.

### 3.5 Fourier Transform Infrared spectrometry (FTIR)

FTIR spectroscopy performs non-destructive analysis of functional groups and chemical bonds. FTIR can distinguish between distinct plant tissues (Barnes et al., 2023). The process of identifying functional groups and characteristic peak values of numerous chemicals components present in the sample offers valuable insights into the chemical composition of plants. The biochemical makeup of plant biomass, including polysaccharides, proteins, carbon, and nitrogen, can be ascertained using FTIR analysis. The principle of Fourier Transform Infrared Spectroscopy (FTIR) works by exposing a sample to infrared light, causing its chemical bonds top absorption specific wavelengths and vibrate uniquely (Principle, 2022). This allows for the identification of chemical compositions and structures. FTIR spectra provide data “absorption versus wavenumber” or “transmission versus wavenumber,” with 3 regions: the far-IR region (<400 cm^-1^), the mid-IR region (400–4000 cm^-1^), and the near-IR region (4,000–13000 cm^-1^) (Nandiyanto et al., 2019).

The FTIR analysis of *Desmodium gangeticum* revealed the presence of various functional groups across its leaves, stems, and roots, indicative of its diverse chemical composition (Fig. 12; Supplementary table 3). Wavenumbers beyond the 3400 cm^-1^ exhibited -OH (Hydroxyl group) suggesting the presence of compounds such as alcohols and phenols (InstaNANO, 2024). Specific peaks at 3417.93, 3422.03, 3696.71, and 3421.05 cm^-1^ indicate O-H stretching vibrations associated with flavonoids and polyphenols (Kale and Dhabe, 2023). Wavenumbers in the range of 2970–2860 cm^-1^ correspond to methyl C-H asymmetric/symmetric stretching, characteristic of alkene and alkyl groups (Nandiyanto et al., 2019; Magdum et al., 2024; Pharmawati and Wrasiati, 2020).

**Fig. 12.**
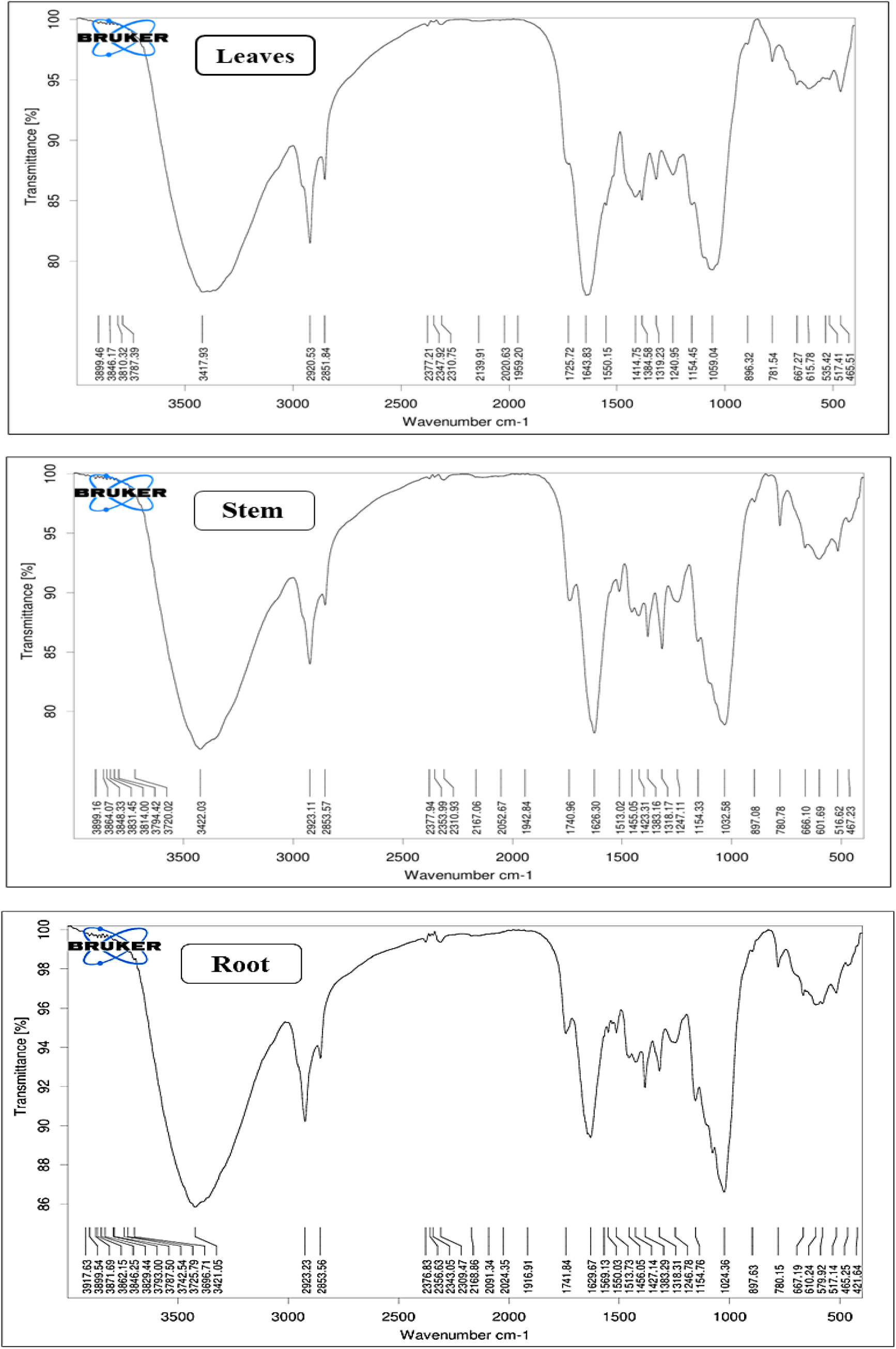
Comparative FTIR Analysis of Leaves, Stem, and Root of *D. gangeticum*.

The presence of O=C=O stretching (Carbonyl Compounds) and CO_2_ groups has been observed in the 2309–2377 cm^-1^ range (Magdum et al., 2024; Pharmawati and Wrasiati, 2020). Wavenumbers between 2140–2175 cm^-1^ indicate S-C≡N stretching (Nandiyanto et al., 2019), while 1940–2140 cm^-1^ showed N=C=S stretching vibrations, indicative of thiocyanate and isothiocyanate groups (Pharmawati and Wrasiati, 2020). The 2024 cm^-1^ wavenumber exhibits C=C=N stretching vibrations (Nandiyanto et al., 2019), suggesting the presence of aromatic ethers and ketenimine. Peaks at 1725 cm^-1^ indicate carboxylic acid groups, while the 1700–1725 cm^-1^ range suggests the presence of carbonyl compounds. The 1741.84 cm^-1^ peak indicates an alkyl carbonate group (1740–1760 cm^-1^). The range of 1650–2000 cm^-1^ shows C-O groups with amide (I/II) vibrations (Magdum et al., 2024; Dalavi and Patil, 2016).

Primary amines with NH bending vibrations have been identified in the 1590–1650 cm^-1^ range, while secondary amines with N-H deformation vibrations have been observed in the 1490– 1580 cm^-1^ range. Aromatic ethers with aryl-O stretching vibrations have been detected in the 1230– 1270 cm^-1^ range (Nandiyanto et al., 2019). The presence of phenol or tertiary alcohol has been indicated in the 1310–1410 cm^-1^ range. Peaks between 700–800 cm^-1^ represented aliphatic halo compounds with C-Cl stretching, and ranges of 600–700 cm^-1^ and 500–600 cm^-1^ indicate C-Br and C-I stretching, respectively. Wavenumbers in the range of 430–500 cm^-1^ indicate the presence of thiols with aryl disulphide (S–S) stretching vibrations (Nandiyanto et al., 2019).

The FTIR spectroscopic analysis of *D. gangeticum* highlights the presence of a diverse array of functional groups, including alkenes, amines, amides, alcohols, phenols, aromatics, carboxylic acids, esters, ethers, organic halogen compounds, and sulphur derivatives. These findings provide insights into the plant’s chemical composition, suggesting its potential medicinal properties attributed to these bioactive constituents. Findings from the present investigation suggest that roots have shown the maximum number of functional groups, followed by leaves and stems. FTIR spectroscopy proves to be a valuable tool for rapid and cost-effective identification of functional groups, aiding in the comparison and prediction of phytoconstituents across medicinal plants.

### 3.6 GC-MS/MS analysis

GC-MS/MS, a powerful analytical technique combines the separation capabilities of gas chromatography with the detection and identification power of mass spectrometry. It allows detailed characterisation of complex mixtures by separating individual components and providing their molecular identification. Gas Chromatography (GC) separates compounds based on volatility and interactions with the stationary phase, using retention time (RT) for identification and quantification. Mass Spectrometry (MS) identifies compounds by their mass-to-charge ratio (m/z). Together, GC and MS offer detailed structural information and confirm compound identities (Vivekanandan-Giri et al., 2011; Gomathi et al., 2013). The methanol extracts of *D. gangetcium* have been analysed by GC-MS/MS. The relative retention times (Rt) and mass spectra of the extract components were compared with the data library. *D. gangeticum* contains volatile chemical compounds in leaves (25) and roots (21) (Supplementary tables 4, 5 and 6; Figs. 13 and 14), belonging to fatty acids, ketones, hydrocarbons, alcohols, esters, etc., that were identified in GC-MS/MS analysis. Both root and leaf extracts have possessed common bioactive compounds, indicating potential for shared therapeutic properties across plant parts. The leaves and root of *D. gangetcium* were found to have a wide range of bioactive phytoconstituents, including tannins, glycosides, alkaloids, flavonoids, steroids, and others, based on the results of GC-MS/MS.

**Fig. 13.**
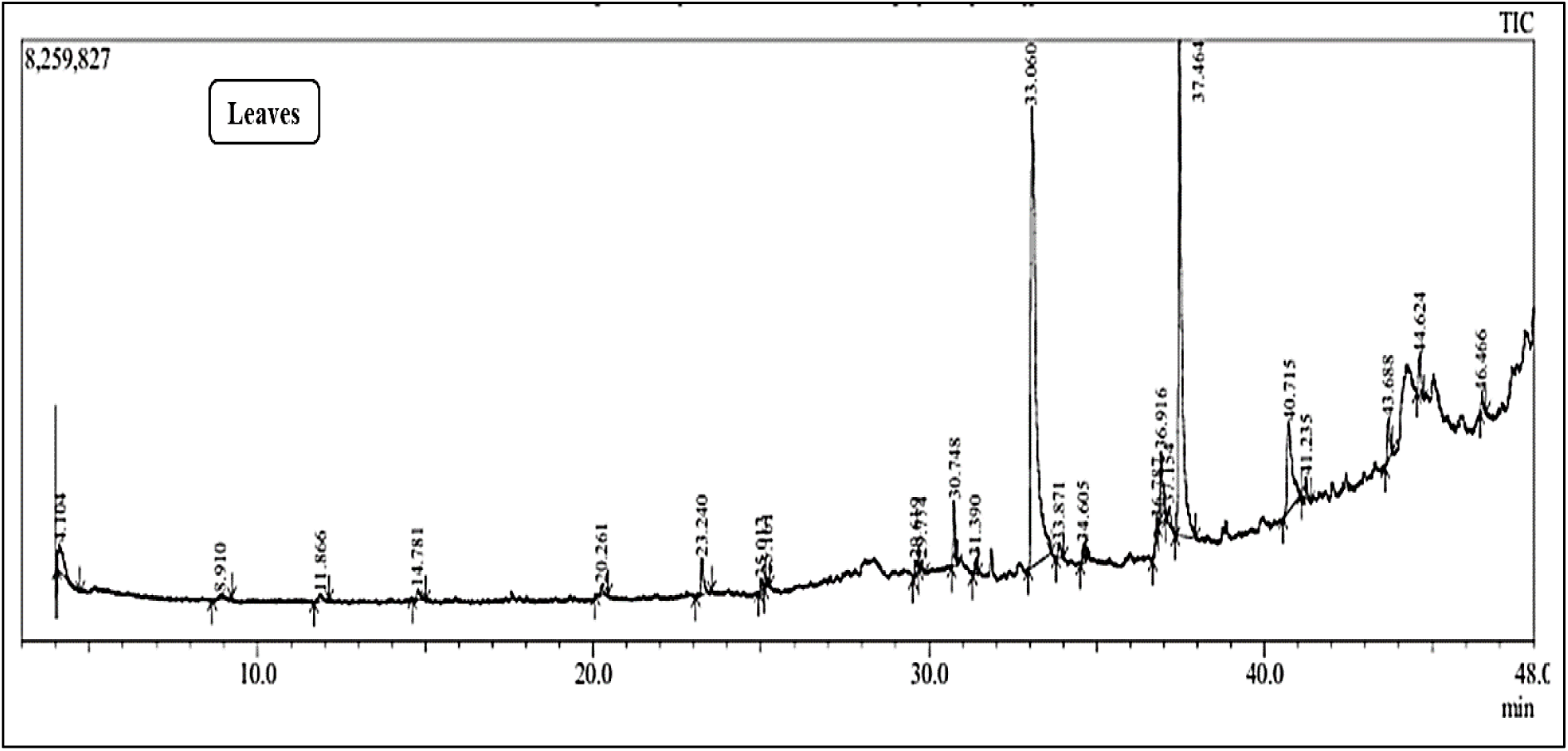
GC-MS/MS Chromatogram of methanolic leaves extract of *D. gangeticum*.

**Fig. 14.**
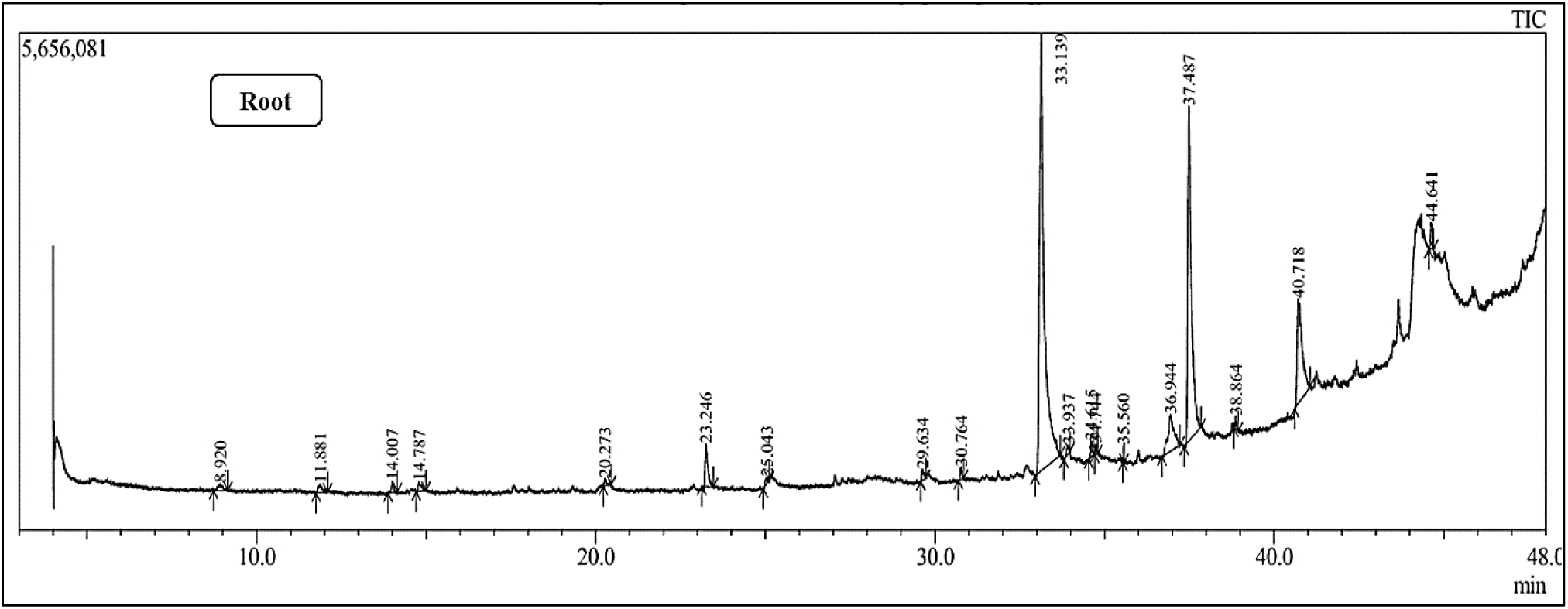
GC-MS/MS Chromatogram of methanolic root extract of *D. gangeticum*.

Many compounds, such as Hexadecanoic acid, methyl ester, and 2,4-Di-tert-butylphenol, exhibit significant anti-inflammatory and antioxidant properties. These activities are essential in preventing oxidative stress and inflammation-related diseases, such as cancer and cardiovascular diseases. Methyl ester, also referred to as methyl hexadecanoate or methyl palmitate, is a lipid mediator with neuroprotective, anti-inflammatory (Aparna et al., 2012), anti-neuroinflammatory, analgesic, and hepatoprotective characteristics. It may be used to treat cancer, cardiometabolic disorders, and rheumatoid arthritis symptoms (Carta et al., 2017; Mancini et al., 2015). Along with these, other compounds exhibit antioxidant and anti-inflammatory activities, which are shown in Supplementary table 4. The compound Bis(2-ethylhexyl) phthalate is antibacterial and larvicidal. It is the precursor to sex hormones like estrogen and progesterone. Trichloroacetic acid, a strong oxidising agent, is disinfectant and antimicrobial agent. Squalene is a biochemical precursor to both steroids and hopanoids (Bloch, 1983). Squalene converted into lanosterol, is used as precursor for steroids such as cholesterol and ergosterol. (Micera et al., 2020; Cerqueira et al., 2016; Zandee 1964). Gas chromatography-mass spectrometry (GC-MS) and mass spectrometry (MS/MS) analyses revealed the presence of numerous bioactive compounds with significant potential against a variety of biological activities. These compounds exhibited antimicrobial, antioxidant, anti-inflammatory, and anticancer properties, indicating their possible applications in pharmaceutical and therapeutic fields.

### 3.6 UPLC-Q-TOF-MS analysis

The UPLC-Q-TOF-MS analysis of leaves and roots of *D. gangeticum* showed diverse bioactive compounds. Excluding unknown compounds (287), 78 of them have been identified as bioactives from leaves of *D. gangeticum* (Supplementary tables 7 and 8: Fig. 15). Most of them are known to be used in drug development, cosmetics, and therapeutics. The identified compounds exhibit significant antioxidant, anti-inflammatory, antimicrobial, anticancer, neuroprotective, antiaging, antiviral, anti-diabetic, antimalarial, analgesic, and antipyretic properties, suggesting their potential in various biomedical applications. While *D. gangeticum* roots showed presence of 192 bioactive compounds (Supplementary tables 7 and 9; Fig. 16).

**Fig. 15.**
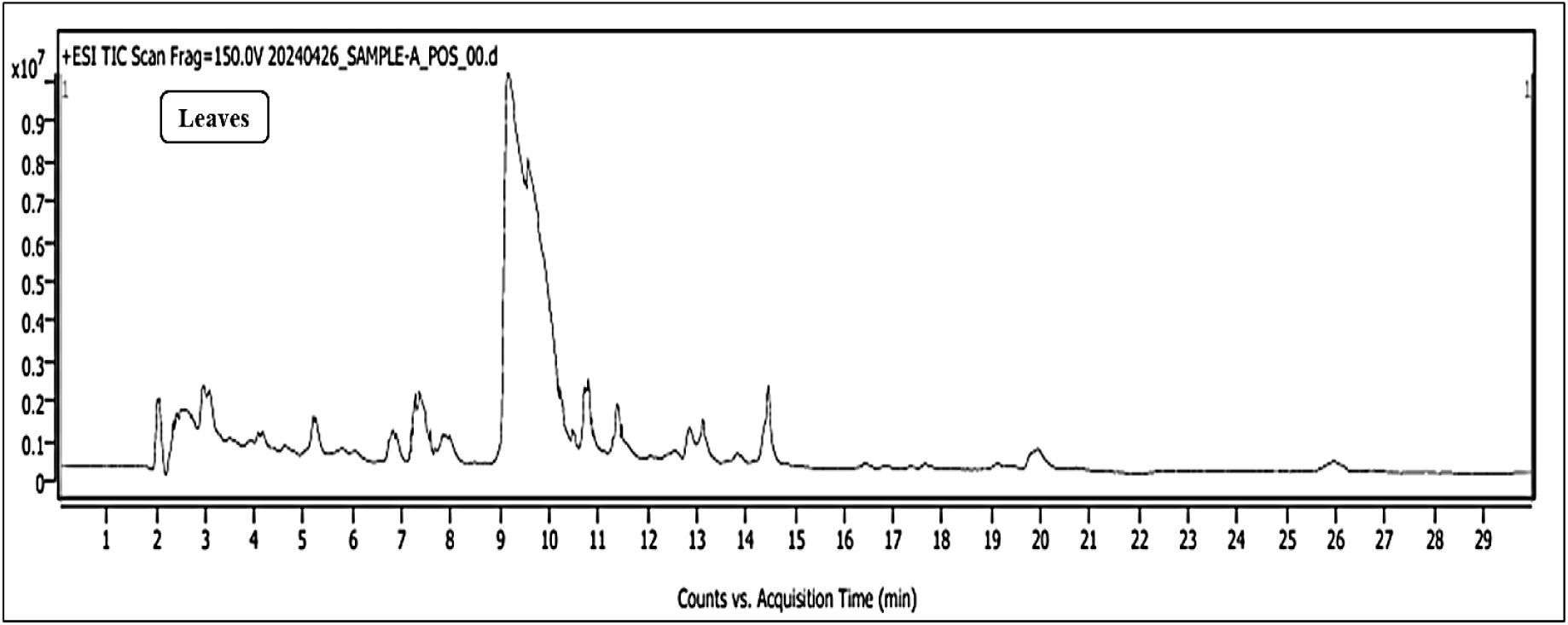
UPLC-Q-TOF-MS Chromatogram of aqueous leaves extracts of *D. gangeticum*.

**Fig. 16.**
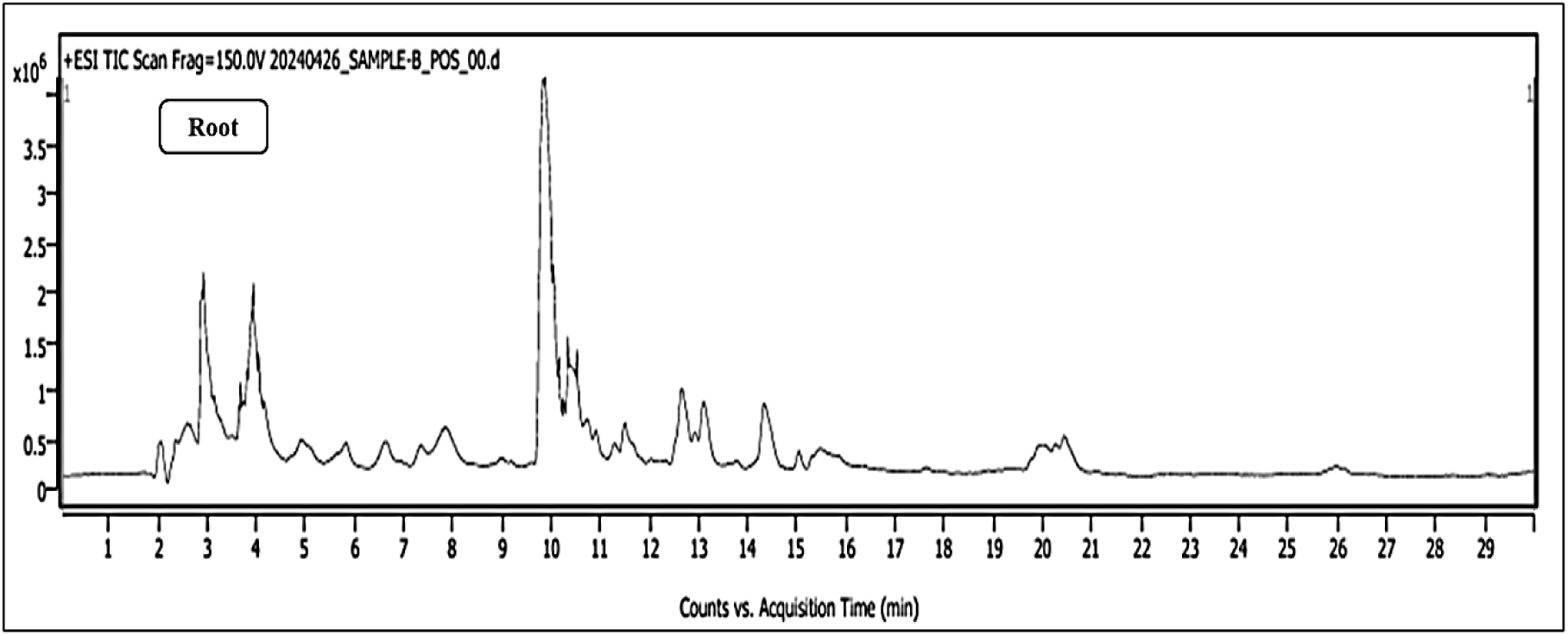
UPLC-Q-TOF-MS Chromatogram of aqueous root extracts of *D. gangeticum*.

## Conclusion

The study reported that the phytochemical screening of different plant parts of *D. gangeticum* with respect to fresh and dried plant parts revealed the presence of bioactive metabolites in leaves, stems and roots. The dried plant parts extracted in different solvents gave promising results than fresh plant parts. Distilled water extracts outperformed methanol and ethanol in terms of phytoconstituent extraction among the three solvents. The other portions of this plant may potentially offer enormous biological potential for the development of several additional herbal formulations to cure ailments, since the roots have previously been utilised to formulate the well-known medicine Dashmularishta. As a result, *D. gangeticum* has great promise as a multifunctional therapeutic agent, and more clinical research should be done to validate its effectiveness. The study also evaluated leaves and stem as other plant parts. While comparing GC-MS/MS and UPLC-Q-TOF-MS analysis many common compounds were found in both. Thus, it might be implied that the entire plant includes therapeutically important compounds. The antibacterial activity against *Staphylococcus aureus* and *Escherichia coli,* underscore the potential of *D. gangeticum* as a natural antimicrobial agent. *D. gangeticum* extract might interact similarly to NSAIDs by demonstrating hypotonicity mediated hemolysis of HRBCs indicating to their anti-inflammatory abilities. The antioxidant, anti-inflammatory and capacity to inhibit bacterial growth make *D. gangeticum* promising for biomedical application.

In conclusion, the results of this study provide valuable understandings into the phytochemical composition and pharmacological activities of *D. gangeticum*. The abundance of phenolics, flavonoids and alkaloids underscore its therapeutic potential. Further studies focussing on identifying and quantifying specific bioactive compound responsible for biological activities would provide different understanding of *D. gangeticum* as a natural antioxidant source.

## Acknowledgment

MAJ would like to acknowledge the financial support received from the SARTHI (Chhatrapati Shahu Maharaj National Research Fellowship-CSMNRF-21). The authors acknowledge the ‘Sophisticated Analytical Instrument Facility (SAIF)’ at Shivaji University, Kolhapur, Maharashtra that accessed *via* I-STEM. The authors would like to acknowledge the Head, Department of Botany, Shivaji University, Kolhapur, India, for providing the excellent laboratory facility. MAJ would like to thank Mr. Gaurav for collection of plant and his support.

**We have used AI for editing and refining the scientific language in the manuscript preparation process.**

---

Supplementary table 1: Antibacterial activity of *D. gangeticum* against S*taphylococcus aureus*.

Supplementary table 2: Antibacterial activity of *D. gangeticum* against *Escherichia coli*.

Supplementary table 3: FTIR analysis of *D. gangeticum* leaves, stem and root.

Supplementary table 4: Bioactive compounds found both in leaf and root extracts of *D. gangeticum*

Supplementary table 5: Bioactive compounds found in leaf extracts of *D. gangeticum*

Supplementary table 6: Bioactive compounds found in root extracts of *D. gangeticum*

Supplementary table 7: Identified bioactive compounds from leaf and root of *D. gangeticum* by UPLC-Q-TOF-MS analysis.

Supplementary table 8: Identified bioactive compounds from leaf of *D. gangeticum* by UPLC-Q-TOF-MS analysis.

Supplementary table 9: Identified bioactive compounds from root of *D. gangeticum* by UPLC-Q-TOF-MS analysis.

