## Supplementary tables for "Chemo-Profiling by UPLC-QTOF-MS, GC-MS/MS analysis and *In Vitro* Bioactivity Assessment of *Desmodium gangeticum* DC"

**Supplementary table 1: Antibacterial activity of *D. gangeticum* against S*taphylococcus aureus.***

| *Sample/ Extracts | Zone of inhibition in 'mm' | | | | |
|---|---|---|---|---|---|
| | **6.25 mg/ml** | **12.5 mg/ml** | **25 mg/ml** | **50 mg/ml** | **100 mg/ml** |
| **Leaf Fresh** | | | | | |
| DGFML | 6±0.1**0.1** | 6±0.2**±0.2** | 7±0.4 | 7±0.2 | 6.±0.8 |
| DGFEL | 7±0.8 | 7±0.6 | 7±0.3 | 7.25±1.2 | 7.75±0.6 |
| DGFDL | 6.75±0.2 | 6.75±0.2 | 7±0.2 | 7.75±0.2 | 9.5±0.4 |
| **Leaf Dry** | | | | | |
| DGDML | 6.5±0.11 | 6.75±1.1 | 6.5±0.2 | 6.5±0.7 | 7.25±0.4 |
| DGDEL | 8±0.2 | 8±0.5 | 7.75±0.16 | 7.25±0.11 | 8.25±0.2 |
| DGDDL | 7.5±0.5 | 6.75±0.4 | 7.25±0.8 | 8±0.3 | **11.5±0.9** |
| **Stem Fresh** | | | | | |
| DGFMS | 7±0.4 | 7.25±0.6 | 7.5±0.9 | 6.75±0.1 | 7±1.4 |
| DGFES | 7±0.7 | 7±0.4 | 6.75±0.15 | 6.75±0.8 | 7.25±0.9 |
| DGFDS | 6.5±0.6**.6** | 6.75±0.9 | 7.25±0.4 | 7.75±0.3 | 9.5±0.6 |
| **Stem Dry** | | | | | |
| DGDMS | 6.5±0.1 | 6.5±0.4 | 6.5±0.5 | 6.5±0.4 | 6.5±0.4 |
| DGDES | 6.5±0.13 | 6.5±0.6 | 6.5±0.3 | 6.5±0.4 | 6.5±0.6 |
| DGDDS | 6.75±0.5 | 7.5±1.1 | 7.5±0.9 | 8±0.15 | 11.25±0.3 |
| **Root Fresh** | | | | | |
| DGFMR | 6.5±0.4 | 7.25±0.5 | 6.75±0.5 | 6.75±0.9 | 6.75±0.5 |
| DGFER | 7.25±0.6 | 8±0.2 | 7.75±0.02 | 7±0.60.65 | 7±0.8 |
| DGFDR | 6.75±0.1 | 7±0.6 | 7.5±0.4 | 8.25±0.5 | 8.75±0.06 |
| **Root Dry** | | | | | |
| DGDMR | 7±0.3 | 7±0.05 | 7±0.2 | 7.5±0.6 | 7±0.2 |
| DGDER | 6.75±0.2 | 7±0.8 | 8±0.6 | 7±0.4 | 8.75±0.6 |
| DGDDR | 6.5±0.4 | 7±0.3 | 7±0.09 | 8±0.1 | 11±0.2 |
| **Streptomycin** | 14±0.6 | 16±0.4 | 20±0.2 | 23±0.19 | **27±0.1** |

*Extracts of 1. DGFML- *D. gangeticum* fresh methanolic leaf, 2. DGFMS- *D. gangeticum* fresh methanolic stem, 3. DGFMR -*D. gangeticum* fresh methanolic root, 4. DGFEL- *D. gangeticum* fresh ethanolic leaf, 5. DGFES- *D. gangeticum*-fresh ethanolic stem, 6. DGFER- *D. gangeticum* fresh ethanolic root,7. DGFDL- *D. gangeticum* fresh distilled water leaf, 8. DGFDS- *D. gangeticum* fresh distilled water stem, 9. DGFDR- *D. gangeticum* fresh distilled water root, 10. DGDML- *D. gangeticum* dry methanolic leaf, 11. DGDMS- *D. gangeticum* dry methanolic stem,12. DGDMR- *D. gangeticum* dry methanolic root, 13. DGDEL- *D. gangeticum* dry ethanolic leaf, 14. DGDES- *D. gangeticum* dry ethanolic stem, 15. DGDER- *D. gangeticum* dry ethanolic root, 16. DGDDL- *D. gangeticum* dry distilled water leaf, 17. DGDDS- *D. gangeticum* dry distilled water stem, 18. DGDDR- *D. gangeticum* dry distilled water root.

### Supplementary table 2: Antibacterial activity of *D. gangeticum* against *Escherichia coli.*

| *Sample/ Extract | Zone of inhibition in 'mm' | | | | |
|---|---|---|---|---|---|
| | **6.25 mg/ml** | **12.5 mg/ml** | **25 mg/ml** | **50 mg/ml** | **100 mg/ml** |
| **Leaf Fresh** | | | | | |
| DGFML | 6.±0.2 | 6±0.5 | 6±0.12 | 6±0.7 | 6±0.09 |
| DGFEL | 6.5±0.4 | 7.5±0.6 | 7±0.08 | 8.25±0.9 | 7.5±0.9 |
| DGFDL | 6.5±0.6 | 7.25±0.4 | 7.25±0.6 | 7.5±0.5 | **10±0.01** |
| **Leaf Dry** | | | | | |
| DGDML | 6.5±0.05 | 6.5±0.7 | 6.5±0.85 | 6.5±0.6 | 6.5±0.5 |
| DGDEL | 6.5±0.3 | 6.5±0.11 | 6.5±0.23 | 6.5±0.7 | 6.5±0.74 |
| DGDDL | 6.25±0.4 | 6.75±0.62 | 7.25±0.41 | 7.75±0.19 | 9±0.21 |
| **Stem Fresh** | | | | | |
| DGFMS | 6.5±0.33 | 6.5±0.36 | 6.5±0.21 | 6.5±0.25 | 6.5±0.1 |
| DGFES | 6.75±0.54 | 6.75±0.45 | 7.25±0.36 | 6.75±0.9 | 7±0.8 |
| DGFDS | 6.5±0.44 | 7±0.22 | 7.75±0.2 | 8.25±0.16 | 9.25±0.15 |
| **Stem Dry** | | | | | |
| DGDMS | 6.25±0.42 | 6.25±0.6 | 6.25±0.2 | 6.25±0.45 | 6.5±0.5 |
| DGDES | 6±0.07 | 6.25±0.39 | 6.25±0.12 | 6.75±0.13 | 6.5±0.26 |
| DGDDS | 6.25±0.2 | 6.5±0.44 | 6.75±0.58 | 7.5±0.64 | 8.5±0.29 |
| **Root Fresh** | | | | | |
| DGFMR | 6.5±0.45 | 6.5±0.14 | 6.5±0.17 | 7±0.06 | 6.25±0.17 |
| DGFER | 6±0.15 | 6±0.22 | 6.25±0.4 | 6.25±0.22 | 6.5±0.42 |
| DGFDR | 6.5±0.28 | 6.5±0.18 | 6.5±0.32 | 7±0.29 | 7.75±0.54 |
| **Root Dry** | | | | | |
| DGDMR | 6.25±0.1 | 6.25±0.09 | 6.5±0.16 | 6.75±0.22 | 6.5±0.1 |

| | | | | | |
|---|---|---|---|---|---|
| DGDER | 6.25±0.13 | 6.25±0.2 | 6.5±0.27 | 6.5±0.4 | 6.5±0.9 |
| DGDDR | 6±0.2 | 6±0.09 | 6.5±0.6 | 7.25±0.74 | 8.75±0.96 |
| **Streptomycin** | 14.5±0.09 | 17±0.1 | 20±0.08 | 22±0.46 | **26±0.15** |

*Extracts of 1. DGFML- *D. gangeticum* fresh methanolic leaf, 2. DGFMS- *D. gangeticum* fresh methanolic stem, 3. DGFMR -*D. gangeticum* fresh methanolic root, 4. DGFEL- *D. gangeticum* fresh ethanolic leaf, 5. DGFES- *D. gangeticum*-fresh ethanolic stem, 6. DGFER- *D. gangeticum* fresh ethanolic root,7. DGFDL- *D. gangeticum* fresh distilled water leaf, 8. DGFDS- *D. gangeticum* fresh distilled water stem, 9. DGFDR- *D. gangeticum* fresh distilled water root, 10. DGDML- *D. gangeticum* dry methanolic leaf, 11. DGDMS- *D. gangeticum* dry methanolic stem,12. DGDMR- *D. gangeticum* dry methanolic root, 13. DGDEL- *D. gangeticum* dry ethanolic leaf, 14. DGDES- *D. gangeticum* dry ethanolic stem, 15. DGDER- *D. gangeticum* dry ethanolic root, 16. DGDDL-*D. gangeticum* dry distilled water leaf, 17. DGDDS- *D. gangeticum* dry distilled water stem, 18. DGDDR- *D. gangeticum* dry distilled water root.

**Supplementary table 3: FTIR analysis of *D. gangeticum* leaves, stem and root.**

| Sr. No. | Wavenumber $cm^{-1}$ | Class | Functional group | Leaves | Stem | Root |
|---|---|---|---|---|---|---|
| 1 | 3696.71 | Alcohol | O-H stretching | ✓ | - | - |
| 2 | 3421.05 | Alcohol | O-H stretching | ✓ | ✓ | ✓ |
| 3 | 2923.23 | Methylene | C-H asymmetric/symmetric stretch | ✓ | ✓ | ✓ |
| 4 | 2853.56 | Methylene | C-H asymmetric/symmetric stretch | ✓ | ✓ | ✓ |
| 5 | 2376.83 | Carbon dioxide | O=C=O stretching | ✓ | ✓ | ✓ |
| 6 | 2356.63 | Carbon dioxide | O=C=O stretching | ✓ | ✓ | ✓ |
| 7 | 2343.05 | Carbon dioxide | O=C=O stretching | - | - | ✓ |
| 8 | 2309.47 | Carbon dioxide | O=C=O stretching | ✓ | ✓ | ✓ |
| 9 | 2168.86 | Thiocyanate | S-C≡N stretching | - | ✓ | ✓ |
| 10 | 2091.34 | Isothiocyanate | N=C=S stretching | - | ✓ | ✓ |
| 11 | 2024.35 | Ketenimine | C=C=N stretching | - | - | ✓ |
| 12 | 2020.63 | Common inorganic ions | Cyanide ion, thiocyanate ion, and related ions | ✓ | - | - |
| 13 | 1916.91 | Aromatic compound | C-H bending | - | - | ✓ |
| 14 | 1741.84 | Carbonyl compound | Alkyl carbonate | - | ✓ | ✓ |
| 15 | 1643.83 | Amide (I/II) | C=O stretch | ✓ | - | - |
| 16 | 1629.67 | Alkene | C=C stretching | - | ✓ | ✓ |
| 17 | 1569.13 | Pri. Amino, Cyclic | Primary amine, NH bend | - | - | ✓ |
| 18 | 1550.03 | N-O stretching | Nitro compound | ✓ | - | ✓ |
| 19 | 1513.73 | N-O stretching | Nitro compound | - | ✓ | ✓ |
| 20 | 1456.05 | R-CH3 alkane | C-H deformation vibration | - | ✓ | ✓ |
| 21 | 1427.14 | Carboxylic acid | O-H bending | - | ✓ | ✓ |
| 22 | 1414.75 | Olefinic alkene | Vinyl C-H in plane bend | ✓ | - | - |
| 23 | 1383.29 | Alcohol and phenoxy compound | Phenol or tertiary alcohol, OH bend | ✓ | ✓ | ✓ |
| 24 | 1318.31 | Alcohol and phenoxy compound | Phenol or tertiary alcohol, OH bend | ✓ | ✓ | ✓ |
| 25 | 1246.78 | Ether and -oxy compounds | Aromatic ethers, aryl -O stretch | ✓ | ✓ | ✓ |
| 26 | 1154.76 | Sec. Amine | C-N stretching | ✓ | ✓ | ✓ |
| 27 | 1059.04 | Amine | C-N stretching | ✓ | - | - |
| 28 | 1024.36 | Amine | C-N stretching | - | ✓ | ✓ |
| 29 | 897.63 | Aromatic group (Aryl) | 1,3 disubstitution (meta) | ✓ | ✓ | ✓ |

| | | | | | | |
|---|---|---|---|---|---|---|
| 30 | 780.15 | Aliphatic halo compound | Aliphatic chloro compounds, C-Cl stretch | ✓ | ✓ | ✓ |
| 31 | 667.19 | Aliphatic organohalo compound | Aliphatic bromo compounds, C-Br stretch | ✓ | ✓ | ✓ |
| 32 | 610.24 | Aliphatic organohalo compound | Aliphatic bromo compounds, C-Br stretch | ✓ | ✓ | ✓ |
| 33 | 579.92 | Aliphatic organohalo compounds | Aliphatic iodo compounds, C-I stretch | - | ✓ | ✓ |
| 34 | 517.14 | Aliphatic organohalo compounds | Aliphatic iodo compounds, C-I stretch | ✓ | ✓ | ✓ |
| 35 | 487.23 | Thiols | Aryl disulfides (S-S stretch) | - | ✓ | - |
| 36 | 465.25 | Thiols | Aryl disulfides (S-S stretch) | ✓ | - | ✓ |
| 37 | 421.64 | Thiols | Aryl disulfides (S-S stretch) | - | - | ✓ |

***(✓)** Presence of Function group**; (-)** Absence of Functional group

**Supplementary table 4: Bioactive compounds found both in leaf and root extracts of *D. gangeticum***

| Sr. No | Name of compound | Functional group/Class | Structural formula | Area % | RT | Biological use | Reference |
|---|---|---|---|---|---|---|---|
| 1 | Undecane | Liquid alkane hydrocarbon | $CH_3(CH_2)_9CH_3$ | 0.34 | 11.86 | Anti-inflammatory, immune suppression, antimutagenic, anti-plasmodial, antiviral | Qin et al., (2019) |
| 2 | Dodecane | Aliphatic compound | $C_{12}H_{26}$ | 0.36 | 14.78 | Antibacterial, antifungal | Padma et al., (2019) |
| 3 | 2,4-Di-tert-butylphenol | Hydroxyl group (OH) attached to the benzene ring | $C_{14}H_{22}O$ | 23.24 | 1.45 | Antioxidant activity, anti-inflammatory activity, cytotoxicity, antifungal | Varsha et al., (2015) |
| 4 | Hexadecane | Alkane hydrocarbon | $C_{16}H_{34}$ | 25.16 | 0.30 | Antibacterial, antioxidant | Yogeswari et al., (2012) |
| 5 | 1-Nonadecene | Alkene | $CH_3(CH_2)_{16}CH=CH_2CH_3(CH_2)_{16}CH=CH_2$ | 29.61 | 0.50 | Antioxidant and anti-inflammatory | Albratty et al., (2023) |
| 6 | Neophytadiene | Diterpene | $C_{20}H_{38}$ | 30.74 | 1.89 | Anti-inflammatory, anticonvulsant | Bhardwaj et al., (2020),<br>Gonzalez-Rivera et al., (2023) |
| 7 | Hexadecanoic acid, methyl ester (methyl palmitate or methyl hexadecanoate) | Fatty acid methyl ester | $CH_3(CH_2)_{14}COOH$ | 33.06 | 40.25 | Anti-inflammatory, neuroprotective, anti-neuroinflammatory, analgesic, antibacterial, hepatoprotective, antirheumatic, anticancer, anti-cardiometabolic | Carta et al., (2017),<br>Aparna et al., (2012)<br>Mancini et al., (2015b) |

| | | | | | | | |
|---|---|---|---|---|---|---|---|
| 8 | Phthalic acid, butyl hexyl ester | Aromatic dicarboxylic acid | HO(O)C−$C_6H_4$−C(O)OH | 33.87 | 0.61 | Antioxidant, antimicrobial, insecticidal | Huang L. et al., (2021) |
| 9 | 9,12-Octadecadienoic acid (Z,Z)-, methyl ester | Fatty acid ester | $CH_3OOC(CH_2)_7CH=CH(CH_2)_7CH=CH(CH_2)_7COOCH_3$ | 36.78 | 0.42 | Anti-inflammatory, anticancer, antimicrobial, antioxidant | Krishnamoorthy and Subramaniam (2014),<br>Rossellia et al., (2007) |
| 10 | 6-Octadecenoic acid, methyl ester, (Z)- | Fatty acid ester | $C_{19}H_{36}O_2$ | 36.91 | 3.12 | Antioxidant, antimicrobial, anticancer | Adegoke et al., (2019) |
| 11 | Methyl stearate | Fatty acid methyl ester (FAMEs) | $C_{19}H_{38}O_2$ | 37.46 | 27.91 | Anti-inflammatory, nematicidal, antinociceptive, antioxidant, antifungal | Adnan et al., (2019) |
| 12 | 9-Octadecenoic acid, 12-hydroxy-, methyl ester | Ester | $C_{19}H_{36}O_3$ | 40.71 | 7.14 | Anti-inflammatory, anticancer,<br>Hypocholesterolemic, dermatitigenic effects | Natarajan et al., (2019),<br>NCBI PubChem 2024 |
| 13 | Methyl 18-methylnonadecanoate | Fatty acid ester | $C_{21}H_{42}O_2$ | 41.23 | 0.47 | Antioxidant, antimicrobial, anticancer | Smolecule |
| 14 | Hexadecanoic acid, (2,2-dimethyl-1,3-dioxolan-4-yl) methyl ester<br>23 (or palmitate ester) | Fatty acid methyl esters | $C_{17}H_{34}O_2$ | 43.68 | 1.88 | Anti-arthritic,<br>anti-inflammatory,<br>anti-ulcer, anti-diuretic,<br>hepatoprotective, neuroprotective | Olivia et al., (2021) |
| 15 | Bis(2-ethylhexyl) phthalate | Ester | $C_{24}H_{38}O_4$ | 44.62 | 1.84 | Antibacterial, larvicidal<br>anti-aging effects | Javed et al., (2022) |

**Supplementary table 5: Bioactive compounds found in leaf extracts of *D. gangeticum***

| Sr. No | Name of compound | Functional group/Class | Structural formula | Area % | | RT | Biological use | Reference |
|---|---|---|---|---|---|---|---|---|
| 1 | 3-Hexadecanol | Secondary hydroxyl group | $C_{16}H_{34}O$ | 1.98 | | 4.10 | Bacterial and plant metabolite | NCBI PubChem 2024 |
| 2 | Tetradecane | Alkane | $CH_3(CH_2)_{12}CH_3$ | 0.21 | | 20.26 | Plant-insect interactions, Biomarker in dog hair to detect visceral leishmaniasis | Yin et al., (2022), |
| 3 | Trichloroacetic acid, undecyl ester | Saturated aliphatic hydrocarbon | $C_{12}H_{21}Cl_3O_2$ | 25.01 | | 0.37 | Antimicrobial, Disinfectant | Karthik et al., (2023) |
| 4 | Heneicosane | Alkane hydrocarbony | $CH_3(CH_2)_{19}CH_3$ | 29.77 | | 0.32 | Antibacterial, Treatment on Gonorrhoea, Syphilis, Tuberculosis | Vanitha et al., (2020), Okechukwu (2020) |
| 5 | 3,7,11,15-Tetramethyl-2-hexadecen-1-ol (Phytol) | Acyclic diterpene alcohol | $C_{20}H_{40}O$ | 31.39 | | 0.43 | Antioxidant, anti-inflammatory, anticancer, antimicrobial, antifungal Fragrance in cosmetic industry | Islam et al., (2018) |
| 6 | Phytol | Diterpene alcohol | $C_{20}H_{40}O$ | 37.15 | | 0.56 | Anticancer, insecticidal, antioxidant, anti-inflammatory, antimicrobial, antifungal, anxiolytic, anticonvulsant, used in the cosmetics industry | Gliszczyńska et al., (2021), Islam et al., (2018), |
| 7 | Methyl 20-methyl-docosanoate (behenic acid methyl ester) | Fatty acid methyl ester | $CH_3(CH_2)_{20}COOCH_3$ | 46.46 | | 0.58 | Anticancer, anti-inflammatory | MedChem Express 2024 |

| | | | | | | | | |
|---|---|---|---|---|---|---|---|---|
| 8 | Tetracosanoic acid, methyl ester (lignoceric acid methyl ester) | Fatty acid methyl ester | $C_{25}H_{50}O_2$ | 48.34 | | 2.55 | Antioxidant, anti-inflammatory | Nyalo et al., (2023) |
| 9 | Stigmasta-3,5-diene | Steroid | $C_{29}H_{48}$ | 48.79 | | 3.36 | Precursor for the synthesis of essential steroids in the body, anti-inflammatory | Smolecule |
| 10 | Supraene | Isoprenoid | $C_{30}H_{50}$ | 49.94 | | 0.49 | Cosmetic industry | Kim and Karadeniz (2012) |

**Supplementary table 6: Bioactive compounds found in root extracts of *D. gangeticum***

| Sr. No | Compound | Functional group/Class | Structural formula | RT | Area % | Biological use | Reference |
|---|---|---|---|---|---|---|---|
| 1 | Cyclohexanol, 2-(1-methylethyl)-(menthol) | 2° alcohol | $C_6H_{11}OH$ | 14.00 | 0.47 | Anticancer, expectorant, biomarkers of tobacco exposure | NCBI PubChem 2024 |
| 2 | Fluoroacetic acid, dodecyl ester | Fluoroacetate ester | $CH_3(CH_2)_{11}CH_2O$ -C(O)-$CH_2F$ | 25.04 | 0.28 | Antifungal, Rodenticide | Hawar et al., (2023), |
| 3 | Heptadecane, 2,6,10,15-tetramethyl- | Sesquiterpenoid | $C_{21}H_{44}$ | 20.27 | 0.34 | Antimicrobial, antioxidant, anti-inflammatory, anticancer | Pena et al., (2019) |
| 4 | Hexadecane, 1-iodo- | | $C_{16}H_{33}I$ | 38.86 | 0.33 | treatment of atopic dermatitis | D. Y. Kim et al., (2022), CHEMI.com |
| 5 | Squalene | Triterpene | $C_{30}H_{50}$ | 49.94 | 2.34 | Antioxidant, anticancer, topical skin lubrication and protection, chemo preventive, precursor to both steroids and hopanoids, cosmetic dermatology, prevention of cutaneous aging | Huang et al., (2009), Micera et al., (2020), Cerqueira et al., (2016), Bloch (1983), Zandee (1964) Lou-Bonafonte et al., (2018) |
| 6 | Tetradecane, 2,6,10-trimethyl- | Saturated alkane hydrocarbon | $C_{17}H_{36}$ | 34.74 | 0.25 | Antimicrobial, antioxidant, antibacterial, antifungal, anticancer | You et al., (2022), Karthik et al., (2023) |

**Supplementary table 7: Identified bioactive compounds from leaf and root of *D. gangeticum* by UPLC-Q-TOF-MS analysis.**

| Sr. No. | Compound name | Chemical formula | Retention Time | Mass |
|---|---|---|---|---|
| | **Phenolics and flavonoids** | | | |
| 1 | Calophyllin B | $C_{18}H_{16}O_4$ | 17.66 | 296.10 |
| 2 | Larixinic Acid | $C_6H_6O_3$ | 7.87 | 126.03 |
| 3 | 4-Prenylresveratrol | $C_{19}H_{20}O_3$ | 10.22 | 296.14 |
| 4 | Fluenetil | $C_{16}H_{15}FO_2$ | 3.33 | 240.14 |
| | **Alkaloids** | | | |
| 1 | Triacanthine | $C_{10}H_{13}N_5$ | 12.61 | 203.11 |
| 2 | N-Methylpelletierine | $C_9H_{17}NO$ | 7.79 | 155.13 |
| 3 | Trachelanthamidine | $C_8H_{15}NO$ | 5.80 | 141.11 |
| 4 | Tranylcypromine | $C_9H_{11}N$ | 11.80 | 133.08 |
| | **Terpenes** | | | |
| 1 | (+/-)-trans- and cis-4,8-Dimethyl-3,7-nonadien-2-ol | $C_{11}H_{20}O$ | 21.58 | 168.15 |
| 2 | (1R,2R,4R)-1,8-Epoxy-p-menthane-2,4-diol | $C_{10}H_{18}O_3$ | 16.43 | 186.12 |
| 3 | 3-Hydroxy-1-methylestra-1,3,5(10),6-tetraen-17-one | $C_{19}H_{22}O_2$ | 10.13 | 282.16 |
| 4 | Gypsogenin | $C_{30}H_{46}O_4$ | 11.60 | 470.34 |
| | **Amino acids** | | | |
| 1 | 1-[(5-Methyl-2-furanyl) methyl] pyrrolidine | $C_{10}H_{15}NO$ | 4.13 | 165.11 |
| 2 | Histamine | $C_5H_9N_3$ | 2.83 | 111.07 |
| 3 | Leucyl-leucine | $C_{12}H_{24}N_2O_3$ | 6.63 | 244.17 |
| 4 | N,N-dimethylhistidine | $C_8H_{13}N_3O_2$ | 11.19 | 183.10 |
| 5 | Phenylalanyl-Methionine | $C_{14}H_{20}N_2O_3S$ | 2.69 | 296.11 |
| 6 | Valinopine | $C_{10}H_{17}NO_6$ | 12.69 | 247.10 |
| 7 | Nalpha-Methylhistidine | $C_7H_{11}N_3O_2$ | 11.06 | 169.08 |
| 8 | Capryloylglycine | $C_{10}H_{19}NO_3$ | 3.87 | 201.13 |
| | **Cyclitols** | | | |
| 1 | (-)-Viburnitol | $C_6H_{12}O_5$ | 3.04 | 164.06 |
| | **Others** | | | |
| 1 | 2-Dodecylbenzenesulfonic acid (DBSA) | $C_{18}H_{30}O_3S$ | 10.80 | 326.18 |
| 2 | 4-Oxocyclohexanecarboxylate | $C_7H_{10}O_3$ | 2.57 | 142.06 |
| 3 | Caffeic aldehyde | $C_{24}H_{28}N_2O_5$ | 16.76 | 424.20 |
| 4 | Methyl oxalate | $C_4H_6O_4$ | 2.66 | 118.02 |

**Supplementary table 8: Identified bioactive compounds from leaf of *D. gangeticum* by UPLC-Q-TOF-MS analysis.**

| Sr. No. | Compound name | Chemical formula | Retention Time | Mass |
|---|---|---|---|---|
| | **Phenolics and flavonoids** | | | |
| 1 | 2,4-Dimethylphenol | $C_8H_{10}O$ | 2.99 | 122.07 |
| 2 | 2-Heptylfuran | $C_{11}H_{18}O$ | 19.33 | 166.13 |
| 3 | 5-Carboxyvanillic acid | $C_9H_8O_6$ | 5.77 | 212.03 |
| 4 | 7-Hydroxy-3-(4-methoxyphenyl)-4-propyl-2H-1-benzopyran-2-one | $C_{19}H_{18}O_4$ | 2.98 | 310.12 |
| 5 | Dihydroferuloylglycine | $C_{12}H_{15}NO_5$ | 18.01 | 253.09 |
| 6 | Haplodimerine | $C_{28}H_{26}N_2O_6$ | 11.94 | 486.18 |
| 7 | m-Cresol | $C_7H_8O$ | 5.23 | 108.05 |
| 8 | N1-Caffeoyl-N10 feruloylspermidine | $C_{26}H_{33}N_3O_6$ | 7.05 | 483.23 |
| 9 | Otobain | $C_{20}H_{20}O_4$ | 9.07 | 324.13 |
| 10 | Tectoridin/phytoestrogen | $C_{22}H_{22}O_{11}$ | 28.91 | 462.11 |
| 11 | Wine lactone | $C_{10}H_{14}O_2$ | 12.39 | 166.09 |
| | **Alkaloids** | | | |
| 1 | 10,16-dihydroxy-palmitic acid | $C_{16}H_{32}O_4$ | 18.35 | 288.22 |
| 2 | 6-hydroxymelatonin | $C_{13}H_{16}N_2O_3$ | 8.82 | 248.11 |
| 3 | Caffeine | $C_8H_{10}N_4O_2$ | 11.62 | 194.08 |
| 4 | Canthin-6-one | $C_{14}H_8N_2O$ | 3.19 | 220.06 |
| 5 | Capsaicin | $C_{18}H_{27}NO_3$ | 16.88 | 305.19 |
| 6 | Hirsuteine | $C_{22}H_{26}N_2O_3$ | 16.05 | 366.19 |
| 7 | Myosmine | $C_9H_{10}N_2$ | 11.77 | 146.08 |
| 8 | Pseudolycorine | $C_{16}H_{19}NO_4$ | 9.00 | 289.13 |
| 9 | Vomicine | $C_{22}H_{24}N_2O_4$ | 16.54 | 380.17 |
| 10 | 4-Guanidino-1-butanol | $C_5H_{13}N_3O$ | 2.97 | 131.10 |
| 11 | Calystegin A3 | $C_7H_{13}NO_3$ | 2.38 | 159.08 |
| 12 | delta-Guanidinovaleric acid | $C_6H_{13}N_3O_2$ | 3.96 | 159.10 |
| 13 | Dioncophylline C | $C_{23}H_{25}NO_3$ | 2.98 | 363.18 |
| | **Terpenes** | | | |
| 1 | Indicumenone | $C_{15}H_{24}O_3$ | 10.80 | 252.17 |
| 2 | Tyromycic acid | $C_{30}H_{44}O_3$ | 11.59 | 452.33 |
| 3 | 1,5-Octadien-3-one | $C_8H_{12}O$ | 10.78 | 124.08 |
| 4 | Estra-1,3,5(10),16-tetraen-3-olbenzoate | $C_{25}H_{26}O_2$ | 16.54 | 358.19 |
| 5 | Stipitatonate | $C_9H_4O_6$ | 14.27 | 207.99 |
| | **Amino acids** | | | |
| 1 | 2-Amino-5-oxohexanoate | $C_6H_{11}NO_3$ | 2.38 | 145.07 |
| 2 | Tyrosyl-Valine | $C_{14}H_{20}N_2O_4$ | 6.81 | 280.14 |
| 3 | (S)-Piperazine-2-carboxamide | $C_5H_{11}N_3O$ | 2.82 | 129.09 |

| | | | | |
|---|---|---|---|---|
| 4 | 5-Methoxytryptophan (5-MTP) | $C_{12}H_{14}N_2O_3$ | 5.09 | 234.10 |
| 5 | Cyclohexylamine | $C_6H_{13}N$ | 4.51 | 99.10 |
| 6 | γ Glutamyl-threonine | $C_9H_{17}N_3O_5$ | 10.84 | 247.11 |
| 7 | Kinetin | $C_{10}H_9N_5O$ | 18.85 | 215.08 |
| 8 | L,L-Cyclo(leucylprolyl) | $C_{11}H_{18}N_2O_2$ | 11.10 | 210.13 |
| 9 | L-isoleucyl-L-proline | $C_{11}H_{20}N_2O_3$ | 3.06 | 228.14 |
| 10 | Tamoxifen N-oxide | $C_{26}H_{29}NO_2$ | 26.14 | 387.22 |
| 11 | N-(1-Deoxy-1-fructosyl) proline | $C_{11}H_{19}NO_7$ | 3.30 | 277.11 |
| 12 | Capryloylglycine | $C_{10}H_{19}NO_3$ | 3.87 | 201.13 |
| 13 | 2 -(Butylamido)-4-hydroxybutanoic acid | $C_8H_{15}NO_4$ | 19.155 | 189.1014 |
| | **Glycosides** | | | |
| 1 | Deslanoside | $C_{47}H_{74}O_{19}$ | 14.26 | 942.48 |
| | **Others** | | | |
| 1 | (E)-2-Butenyl-4-methyl-threonine | C9 H17 N O3 | 3.676 | 187.12 |
| 2 | 2-Methyl-4-heptanone, (methyl isobutyl ketone) | C9 H18 O | 17.800 | 142.13 |
| 3 | 3-(Pyrazol-1-yl)-L-alanine | $C_6H_9N_3O_2$ | 11.53 | 155.06 |
| 4 | 6-(2-Chloroallylthio)purine (6-TG) | $C_8H_7ClN_4S$ | 2.96 | 226.00 |
| 5 | gamma-Nonalactone | $C_9H_{16}O_2$ | 13.91 | 156.11 |
| 6 | L-Phenylglycine | $C_8H_9NO_2$ | 3.66 | 151.06 |
| 7 | Norvaline | $C_5H_{11}NO_2$ | 2.38 | 117.07 |
| 8 | Ribothymidine | $C_{10}H_{14}N_2O_6$ | 12.98 | 258.08 |
| 9 | Z)-6-Nonenal | $C_9H_{16}O$ | 16.43 | 140.12 |
| 10 | γ-Glutamyl ornithine | $C_{10}H_{19}N_3O_5$ | 12.06 | 261.13 |

**Supplementary table 9: Identified bioactive compounds from root of *D. gangeticum* by UPLC-Q-TOF-MS analysis.**

| Sr. No. | Compound name | Chemical formula | RT | Mass |
|---|---|---|---|---|
| | **Phenolics and Flavonoids** | | | |
| 1 | 4,4'-Methylenebis(2,6-di-tert-butylphenol) | $C_{29}H_{44}O_2$ | 11.49 | 424.33 |
| 2 | 4-Benzyloxy-2'-hydroxy-3',4',5',6'-tetramethoxychalcone | $C_{26}H_{26}O_7$ | 14.09 | 450.17 |
| 3 | N-(2,5-Dihydroxyphenyl) pyridinium | $C_{11}H_{10}NO_2$ | 10.44 | 188.06 |
| | **Alkaloids** | | | |
| 1 | Swainsonine | $C_8H_{15}NO_3$ | 2.78 | 173.10 |
| 2 | Vinpocetine | $C_{22}H_{26}N_2O_2$ | 16.78 | 350.20 |
| 3 | Brucine | $C_{23}H_{26}N_2O_4$ | 13.81 | 394.19 |
| 4 | Conhydrine | $C_8H_{17}NO$ | 2.90 | 143.13 |
| 5 | N-Demethylnarwedine | $C_{16}H_{17}NO_3$ | 12.99 | 271.11 |
| 6 | Raucaffrinoline | $C_{21}H_{24}N_2O_3$ | 3.93 | 352.17 |
| 7 | Scopoline | $C_8H_{13}NO_2$ | 7.72 | 155.09 |
| 8 | Acetylpseudotropine | $C_{10}H_{17}NO_2$ | 9.00 | 183.12 |
| | **Terpenes** | | | |
| 1 | 4,4-Difluoropregn-5-ene-3,20-dione | $C_{21}H_{28}F_2O_2$ | 16.78 | 350.20 |
| 2 | Bruceoside A | $C_{32}H_{42}O_{16}$ | 11.861 | 682.25 |
| 3 | Capsianoside V | $C_{26}H_{42}O_{10}$ | 13.34 | 514.27 |
| 4 | Nogalamycin | $C_{39}H_{49}NO_{16}$ | 14.22 | 787.30 |
| | **Amino acid** | | | |
| 1 | DL-2-Aminooctanoic acid | $C_8H_{17}NO_2$ | 5.83 | 159.12 |
| 2 | Altretamine | $C_9H_{18}N_6$ | 19.40 | 210.15 |
| 3 | Argininosuccinic acid (ASA) | $C_{10}H_{18}N_4O_6$ | 11.02 | 290.12 |
| 4 | Moschamine | $C_{20}H_{20}N_2O_4$ | 13.17 | 352.14 |
| | **Glycosides** | | | |
| 1 | 6S,9R-Dihydroxy-4,7E-megastigmadien-3-one9-[apiosyl-(1->6)-glucoside] | $C_{24}H_{38}O_{12}$ | 12.57 | 518.23 |
| 2 | trans-p-Menthane-1,7,8-triol 8-glucoside | $C_{16}H_{30}O_8$ | 9.58 | 350.19 |
| | **Others** | | | |
| 1 | (E)-2-Butenyl-4-methyl-threonine | $C_9H_{17}NO_3$ | 3.55 | 187.12 |
| 2 | N(alpha)-t-Butoxycarbonyl-L-leucine | $C_{11}H_{21}NO_4$ | 4.74 | 231.14 |
| 3 | (+/-)-3-[(2-methyl-3-furyl)thio]-2-butanone | $C_9H_{12}O_2S$ | 2.04 | 184.05 |

| | | | | |
|---|---|---|---|---|
| 4 | 2,3-Dihydroxycyclopentaneundecanoic acid (Polyhydroxyalkanotes PHAs) | $C_{16}H_{30}O_4$ | 17.94 | 286.21 |
| 5 | 3Z-hexenal | $C_6H_{10}O$ | 2.90 | 98.07 |
| 6 | 3Z-hexenol | $C_6H_{12}O$ | 2.96 | 100.08 |
| 7 | 7,8-Dihydroxykynurenate | $C_{10}H_7NO_5$ | 15.59 | 221.03 |
| 8 | Deoxycoformycin | $C_{11}H_{16}N_4O_4$ | 10.42 | 268.11 |
| 9 | Hematoporphyrin | $C_{34}H_{38}N_4O_6$ | 11.97 | 598.28 |
| 10 | L-Homoserine lactone | $C_4H_7NO_2$ | 2.64 | 101.04 |
| 11 | Pheophorbide a | $C_{35}H_{36}N_4O_5$ | 14.13 | 592.26 |
